# Single-base precision design of CRISPR-Cas13b enables systematic silencing of oncogenic fusions

**DOI:** 10.1101/2022.06.22.497105

**Authors:** Wenxin Hu, Amit Kumar, Shijiao Qi, Teresa Sadras, Joshua ML Casan, David Ma, Lauren M Brown, Michelle Haber, Ilia Voskoboinik, Joseph A Trapani, Paul G Ekert, Mohamed Fareh

## Abstract

Precision oncology programs can rapidly identify oncogenic gene fusions in individual patients^1–3^. However, despite their established oncogenic status, the vast majority of gene fusions remain ‘*undruggable*’ due to the lack of specific inhibitory molecules^4, 5^. Here, we establish *Psp*Cas13b, a poorly characterized programmable RNA nuclease, as a versatile tool to silence various oncogenic fusion transcripts. Our <u>Si</u>ngle-<u>B</u>ase <u>Til</u>ed crRNA screens (<u>SiBTil</u>), unbiased computational analysis, and comprehensive spacer-target mutagenesis revealed key determinants of *Psp*Cas13b activity. *De novo* design of crRNAs harbouring basepaired or mismatched guanosine bases at key spacer positions greatly enhances the silencing efficacy of otherwise inefficient crRNAs, expanding the targeting spectrum of this enzyme. We also reveal the interface between mismatch tolerance and intolerance, which unlocks an unexpected single-base precision targeting capability of this RNA nuclease. Notably, our *de novo* design principles enable potent and selective silencing of various gene fusion transcripts and their downstream oncogenic networks, without off-targeting of non-translocated variants that share extensive sequence homology. We demonstrate that *Psp*Cas13b targeting the breakpoint of fusion transcripts enables efficient suppression of ancestral and single-nucleotide mutants (e.g. BCR-ABL1 T315I) that often drive clinical cancer relapse. Collectively, this study provides new design principles for *Psp*Cas13b programming to specifically recognise and degrade any ‘*undruggable’* fusion oncogenic transcript, thus providing a new conceptual framework for personalized oncology.

## MAIN

Recent advances in next-generation sequencing enables rapid identification of oncogenic transcripts in individual patients within actionable time frames^1–3^. However, many of such driver mutations cannot be targeted due to the lack of specific inhibitory molecules^4, 5^. Numerous fusion genes generated by chromosomal translocations demonstrate cogent oncogenic activity, but also remain largely ‘*undruggable*’ with conventional therapeutics^6^. Personalized targeting of these fusion structural variants at the protein level has proven to be challenging^7^, and in the rare cases where small molecule inhibitors are available, treatments often result in rapid development of drug-resistance and disease relapse^8^. Alternatively, the unique chimeric sequence at the breakpoint of the two genes at the transcript level represents a currently unexplored, tractable, and impactful target for sequence-specific silencing with programmable RNA nucleases.

The type VI CRISPR (clustered regularly interspaced short palindromic repeats) effectors termed CRISPR-Cas13 (Cas13) are programmable RNA-guided targeting enzymes that exclusively degrade single-stranded RNAs (ssRNAs) with high efficacy and specificity^9,10^. Recent studies have deployed Cas13 systems in a variety of targeted RNA manipulations including, nucleic-acid detection^11–13^, precise RNA base editing^14, 15^, and viral suppression^16–18^. The efficiency and reversibility of RNA targeting with Cas13 represents a promising modality to specifically edit oncogenes without risking permanent alteration of the genome in somatic and germline cells, an inherent limitation of DNA-editing CRISPR enzymes^19–21^. Therefore, Cas13 is highly attractive for targeting aberrant fusion transcripts that drive various human genetic diseases including cancer. The *Psp*Cas13b ortholog appears to possess high silencing efficiency and specificity, underscoring its suitability for targeted gene silencing in human cells^22^. However, the poor understanding of the molecular principles governing *Psp*Cas13b target recognition and cleavage limits the development of this tool for exclusive targeting of the breakpoint of any fusion transcripts.

### crRNAs silencing efficiency is highly variable

To elucidate *Psp*Cas13b crRNA design principles, we developed a quantitative fluorescence-based silencing assay, in which we targeted the transcript of the mCherry reporter gene. To achieve this, we co-transfected HEK 293T cells with three plasmids encoding mCherry, *Psp*Cas13b-BFP, and either non-targeting (NT) or mCherry-targeting crRNA (**Figure 1a**). Fluorescence microscopy analysis of cells transfected with mCherry targeting crRNAs showed pronounced silencing activity, contrasted with no appreciable silencing in cells expressing NT crRNAs (**Figure 1b**).

**Figure 1.**
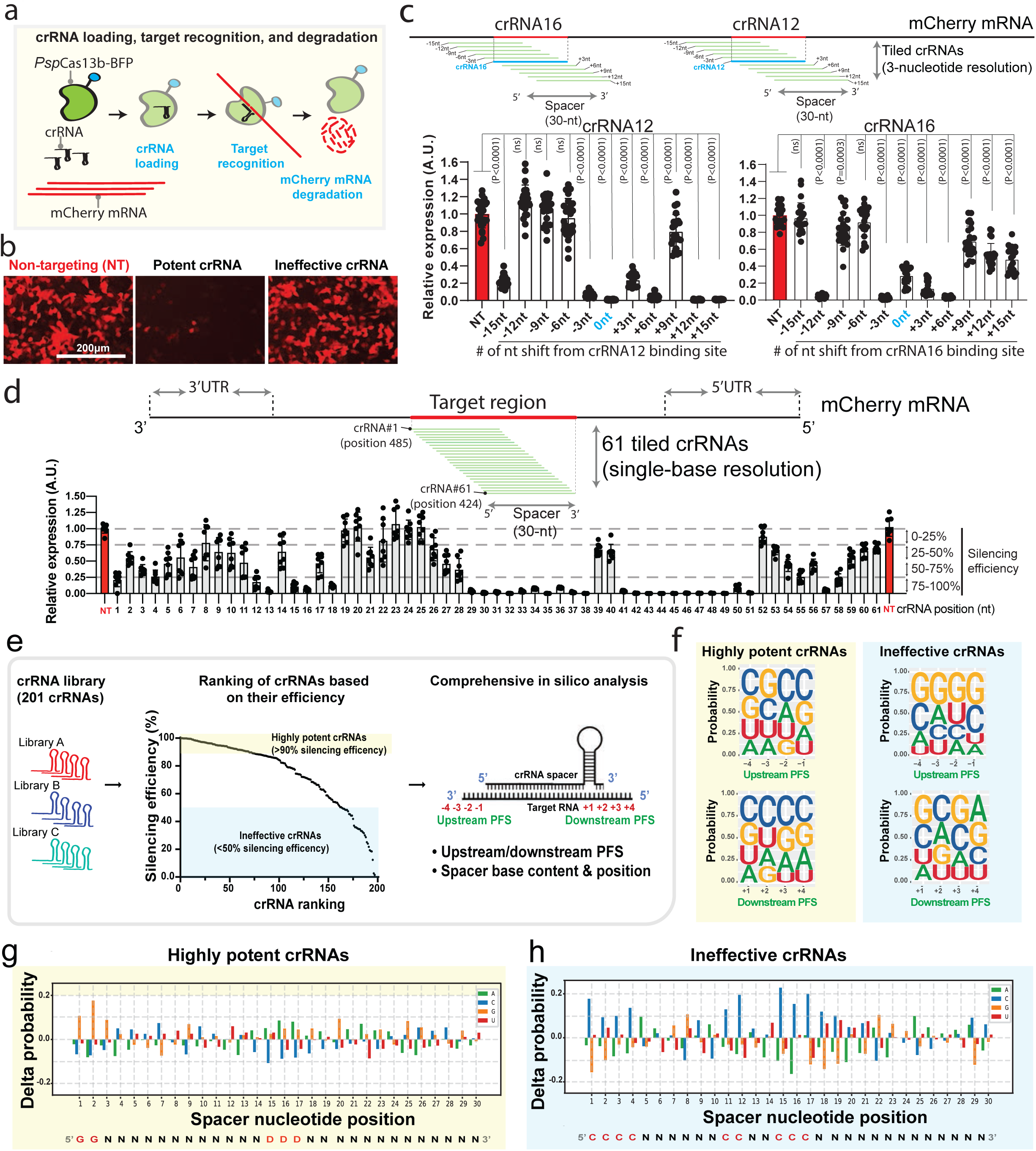
Single-base tiled crRNAs and *in silico* analysis of their silencing profiles revealed key design principles. **(a)** Schematic of *Psp*Cas13b silencing assay used to track the recognition and degradation of mCherry RNA. **(b)** Representative fluorescence microscopy images show the silencing of mCherry transcripts with a potent or ineffective targeting crRNA versus a non-targeting (NT) control crRNA in HEK 293T cells. **(c)** The schematic illustrates mCherry RNA regions covered by 3-nucleotide increments tiled crRNAs. Quantification of silencing efficiency with tiled crRNAs targeting the mCherry regions surrounding crRNA12 (left panel) and crRNA16 (right panel). Data points in the graph are normalized mean fluorescence from 4 representative fields of view per experiment imaged in *N* = 4. The data are represented in arbitrary units (A.U.). Errors are SD and *p*-values of one-way Anova test are indicated (95% confidence interval). **(d)** The schematic shows the sequence of mCherry RNA covered by 61 single-nucleotide resolved tiled crRNAs around the targeted region of crRNA12. The graph quantifies the silencing efficiency obtained with 61 tiled crRNAs in HEK 293T cells. Data points in the graph are normalized mean fluorescence from 4 representative fields of view imaged in *N* = 2. The data are represented in arbitrary units (A.U.). Errors are SD with a 95% confidence interval. *N* is the number of independent biological experiments. Source data are provided as a Source data file. **(e)** Schematic of the bioinformatics pipeline used to investigate various parameters that impact *Psp*Cas13b silencing. PFS positions (4 nucleotides surrounding the 5’ and 3’ end of the targeted region that basepair with the spacer) are indicated. 201 crRNAs are ranked based on their silencing efficiency. The highly potent crRNAs that achieved >90% silencing efficiency (yellow box) and the ineffective crRNAs that achieved <50% silencing efficiency (blue box) are analysed for PFS and spacer nucleotide positions. **(f)** Position Weight Matrices (PWMs) depicting the positional nucleotide probabilities of upstream or downstream PFS in either the highly potent (left panel) or ineffective crRNAs (right panel). **(g-h)** Delta nucleotide probabilities of the highly potent **(g)** and ineffective **(h)** crRNA spacer sequences that compare filtered spacer nucleotide positions to the baseline nucleotide distribution. Spacer nucleotide positions-based formulas to predict highly potent and ineffective spacer sequences. The red colour in the spacer sequence highlights the position of key residues.

Next, we questioned whether parameters such as efficiency of crRNA transcription, crRNA loading, spacer nucleotide composition, target accessibility, and the presence of a potential protospacer-flanking sequence (PFS) may influence the efficiency of *Psp*Cas13b and could lead to variability in the silencing profiles of various crRNAs. We designed 16 crRNAs with spacer sequences that fully basepair with the coding sequence of the mCherry mRNA at various positions (**Extended Data Fig. 1a**). To accurately determine the silencing efficacy of each crRNA in this cohort, we performed crRNA dose-dependent silencing assays in which cells were transfected with 0, 1, 5, and 20 ng of each of 16 mCherry-targeting crRNAs and quantitated silencing efficiency. mCherry targeting crRNAs demonstrated dose-dependent silencing. However, we noticed marked differences in the silencing efficacy of the various crRNAs, even when they are designed to target neighbouring RNA locations (**Figure 1b; Extended Data Fig. 1b-1d**). This finding suggested there are key determinants of *Psp*Cas13b efficacy beyond target accessibility. Identifying such determinants is crucial for efficient reprogramming.

### Single-base tiled crRNAs reveal hidden parameters

To further understand the spectrum of crRNA potency, we investigated the silencing activity of *Psp*Cas13b across a defined targeted region, reasoning that silencing efficiency is likely intrinsic to the spatial characteristics of the crRNA sequence and binding sites. We focused our study on crRNA12 (binding position #455) and crRNA16 (#655) that exhibited high and moderate silencing, respectively (**Extended Data Fig. 1a-1d**). We designed 3-nucleotide resolution tiled crRNAs spanning a 30-nucleotide target region surrounding crRNA12 and crRNA16 binding positions (**Figure 1c; Extended Data Fig. 2a**). In this tiled design, each adjacent crRNAs are spaced by 3 nucleotides, thus silencing profiles should reveal the relationship between efficacy, the sequence of the spacer-target, and target accessibility. We again observed considerable heterogeneity in the potency of these tiled crRNAs despite their physical proximity, with some adjacent crRNAs demonstrating contrasted silencing efficacy. These data indicated that physical barriers such as RNA binding proteins or structured RNA motifs are unlikely to explain the fluctuation in silencing between spatially adjacent crRNAs that are separated by just 3 nucleotides (**Figure 1c; Extended Data Fig. 2a-2c**).

To further enhance our understanding, we maximised the spatial resolution of this approach by designing 61 tiled crRNAs with single-base incremental targeting of the region between nucleotide position 424 and 485 of the mCherry coding sequence (**Figure 1d**). Consistent with previous data, we again observed markedly diverse silencing profiles of neighbouring crRNAs. For instance, crRNA13 achieved silencing exceeding 95% efficiency, but shifting the targeted region by only 1 nucleotide (crRNA14) dramatically reduced efficiency to ∼30%. Similarly, crRNA51 yielded ∼99% silencing efficiency while its adjacent crRNA52 did not show any appreciable silencing activity (**Figure 1d**).

These data strengthen our contention that silencing efficacy is unlikely to be solely dependent on the target accessibility, and that other factors including specific nucleotide positions within the spacer or target, and a possible PFS, may all influence key steps of target silencing such as crRNA transcription, loading, and target recognition.

### *In silico* analysis of 201 crRNAs revealed key design principles

In an effort to uncover fundamental principles that dictate *Psp*Cas13b silencing efficiency, we expanded our dataset by analysing the silencing profiles of 201 individual crRNAs targeting various transcripts^16^. First, we questioned whether the folding of crRNA or target, spacer-target stability, and spacer nucleotide content correlate with pspCas13b potency. The data suggest that the folding of the crRNA and the targeted sequence into complex secondary structures can only moderately limit *Psp*Cas13b silencing efficiency, possibly perturbing crRNA loading or target accessibility (**Extended Data Fig. 3**). Whereas enriched C nucleotides in spacers exhibited a strong negative correlation with crRNA potency (*r*=-0.30; *p*<0.0001) (**Extended Data Fig. 3&4**).

Next, we pooled these 201 crRNAs and ranked them by silencing efficiency. crRNA that achieved >90% silencing efficiency were designated as potent crRNAs and those with less than 50% efficiency were considered ineffective crRNAs. crRNAs with ambiguous silencing profiles (efficiencies ranging from 50 to 90%) were excluded from the analysis. We sought to identify molecular features capable of differentiating potent and ineffective crRNA cohorts (**Figure 1e**). Many CRISPR variants possess an upstream or downstream protospacer flanking sequence (PFS) that restricts targeting activity and prevents degradation of their own nucleic acids^23^. To investigate the existence of a PFS that could constrain *Psp*Cas13b silencing, we generated weight matrix plots that analyse nucleotide composition at each position of four bases upstream and downstream of the targeted sequence in the highly potent and ineffective cohorts of crRNAs. There was no detectable bias in nucleotide composition at various target flanking sites, suggesting that *Psp*Cas13b activity is not subject to PFS motifs in mammalian cells (**Figure 1f**).

Last, we questioned whether the nucleotide composition of the spacer could influence *Psp*Cas13b silencing efficiency. Nucleotide content analysis of the filtered crRNA cohorts revealed an enrichment of G bases in the potent group, and enrichment of C bases in the ineffective crRNA cohort (**Extended Data Fig 4a-4e**), indicating that G-enriched spacers are associated with higher potency, whereas C-enriched spacers are associated with low potency.

To reveal the relevance of G and C bases at specific positions within the spacer sequence, we conducted unbiased analyses of nucleotide composition at all 30 positions of the spacer in highly potent and ineffective crRNA cohorts. We used weight matrix plots and Delta probability analysis to compare spacer nucleotide composition at all positions between filtered and unfiltered samples (**Figure 1g-1h; Extended Data Fig. 2d-2f**), and revealed marked differences in nucleotide positions between the two crRNA cohorts. We show that G bases at the 5’end, particularly a GG sequence at the first and second positions was strongly associated with highly potent crRNAs. Conversely, G nucleotides were depleted and C bases were enriched at the 5’end of spacers in the ineffective crRNA cohort. In addition to this C-rich motif at the 5’end of ineffective crRNAs, we also identified a significant enrichment of C bases at positions 11, 12, 15, 16, and 17 (**Figure 1g-1h; Extended Data Fig. 2d-2f**). These data revealed key nucleotide positions that may determine the potency of crRNAs, which could serve as predictive parameters of crRNA potency.

### Functional validation of crRNA prediction and design

The above *in silico* analysis enabled us to generate a formula to predict potent and ineffective crRNAs. We postulated that potent crRNAs should include GG sequence at the first and second position of the spacer and should lack C bases in position 11, 12, 15, 16, and 17 (***GG***NNNNNNNN***DD***NN***DDD***NNNNNNNNNNNNN; D is a G, U, or A nucleotide; N is any nucleotide). We also hypothesised that crRNAs containing C in spacer positions 1, 2, 3, 4, 11, 12, 15, 16, and 17 are predicted to yield poor silencing efficiency (***CCCC***NNNNNN***CC***NN***CCC***NNNNNNNNNNNNN).

We tested the predictive accuracy of these spacer-based formulas through prospective unbiased design of crRNAs targeting EGFP and TagBFP, two mRNA targets we had not investigated previously. Notably, out of 21 predicted potent crRNAs, 20 achieved very high silencing efficiency of either EGFP or TagBFP mRNA. Conversely, the majority of predicted ineffective crRNAs failed to efficiently silence EGFP and TagBFP transcripts (**Figure 2a-2f**). By formulating our prediction from a pre-existing dataset, and validating its accuracy in heretofore untargeted transcripts, these data demonstrate our spacer nucleotide-based formula to be both accurate and generalisable, and demonstrate its utility in crRNA design for silencing any transcript of interest.

**Figure 2.**
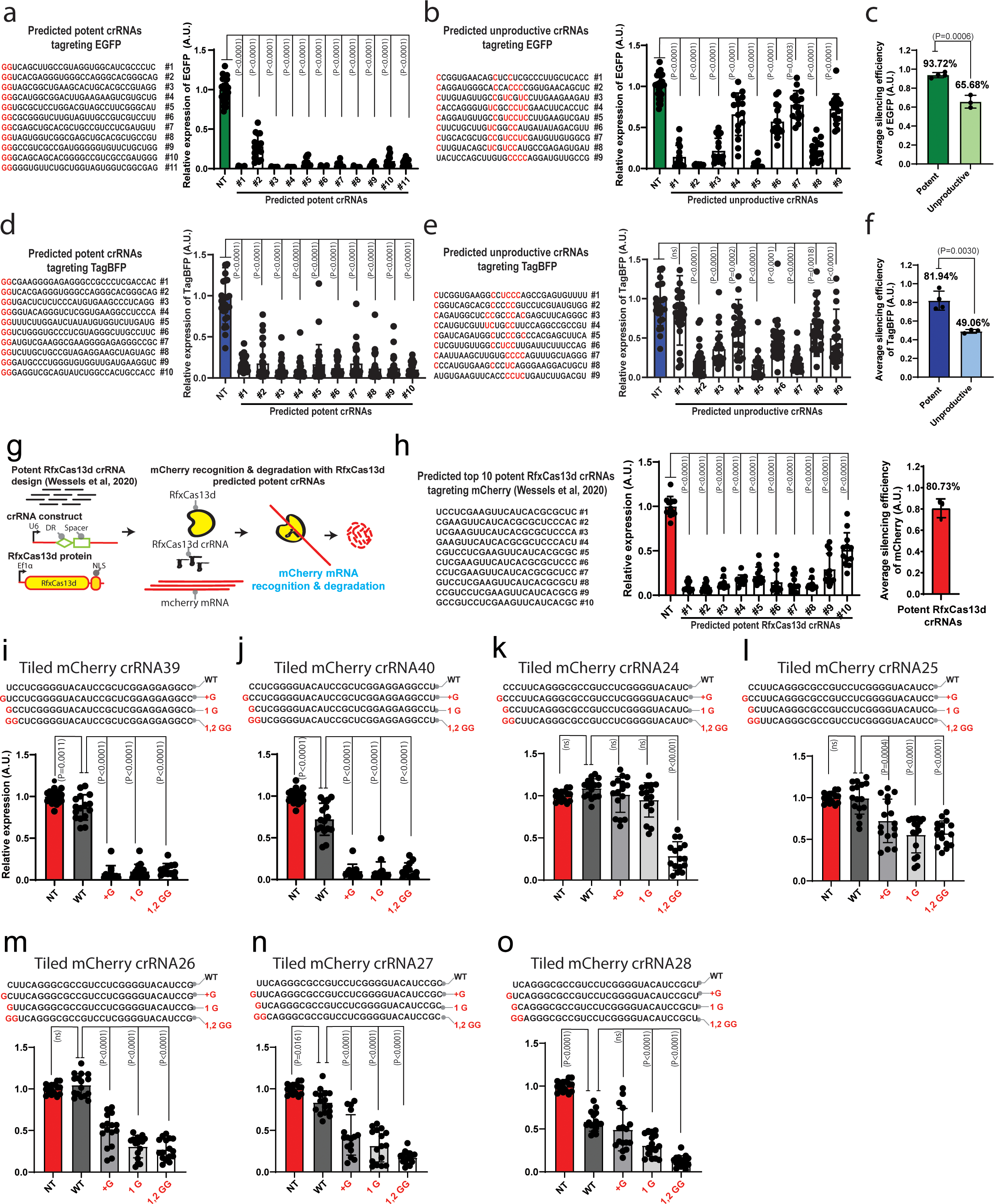
Functional validation of *Psp*Cas13b crRNA prediction and design. **(a-c)** Validation of spacer nucleotide preferences through the targeting EGFP in HEK 293T cells**. (a)** Prospective design and validation of predicted potent crRNAs harbouring a ‘GG’ motif at 5’ end of spacers targeting EGFP transcript. **(b)** Prospective design and validation of predicted ineffective crRNAs lacking 5’ GG motif and harbouring ‘C’ bases at the central region of spacers targeting EGFP transcript. Data points in the graph are mean fluorescence from 4 representative field of views per condition imaged; *N* =3 or 4. The data are represented in arbitrary units (A.U.). Errors are SD and *p*-values of one-way Anova test are indicated (95% confidence interval). **(c)** The average silencing efficiency of predicted potent and ineffective crRNAs targeting EGFP transcripts. *N* = 3 or 4; Data are normalized means and errors are SE; Results are analysed by unpaired two-tailed Student’s t-test (95% confidence interval). **(d-f)** Validation of spacer nucleotide preferences revealed in the screen by targeting TagBFP transcript in HEK 293T cells**. (d)** Prospective design and validation of predicted potent crRNAs harbouring a ‘GG’ motif at 5’ end of spacers targeting TagBFP transcript. **(e)** Prospective design and validation of predicted ineffective crRNAs lacking 5’ GG motif and harbouring ‘C’ bases at the central region of spacers targeting TagBFP transcript. Data points in the graph are mean fluorescence from 4 representative fields of view per condition imaged; *N* =3 or 4. The data are represented in arbitrary units (A.U.). Errors are SD and *p*-values of one-way Anova test are indicated (95% confidence interval). **(f)** The average silencing efficiency of predicted potent and ineffective crRNAs targeting TagBFP transcripts. Data points in the graph represent independent biological replicates. *N* = 3 or 4; Data are normalized means and errors are SE; Results are analysed by unpaired two-tailed Student’s t-test (95% confidence interval). **(g)** Schematic of *Rfx*Cas13d silencing assay to target mCherry transcript in HEK 293T cells using potent crRNAs predicted by the *Rfx*Cas13d guide prediction platform published by Wessels et al^36^. **(h)** The spacer sequences of the top 10 potent *Rfx*Cas13d crRNAs targeting mCherry transcripts predicted with Wessels et al tool^36^ (left). Data points in the graph (middle) are mean fluorescence from 4 representative field of views per condition imaged; *N* =3. The data are represented in arbitrary units (A.U.). Errors are SD and *p*-values of one-way Anova test are indicated (95% confidence interval). The average silencing efficiency of predicted potent *Rfx*Cas13d crRNAs targeting mCherry transcripts (Wessels et al tool^36^) is shown at the right-side graph. Data points in the graph represent independent biological replicates. *N* = 3; Data are normalized means and errors are SE (95% confidence interval). **(i-o)** Incorporation of a G-rich motif at the 5’ end of ineffective spacer sequences targeting mCherry through G-nucleotide insertion or substitution greatly enhanced their silencing efficiency. Data points in the graph are mean fluorescence from 4 representative field of views per condition imaged; *N* =3 or 4. The data are represented in arbitrary units (A.U.). Errors are SD and *p*-values of one-way Anova test are indicated (95% confidence interval). *N* is the number of independent biological replicates. Source data are provided as a Source data file.

Next, we compared the efficiency of our design to the benchmark crRNA design tool that is available for *Rfx*Cas13d (**Figure 2g**). We selected 10 top predicted potent crRNAs for *Rfx*Cas13d targeting mCherry and probed their silencing efficiency, which achieved an average silencing of 80.7% (**Figure 2h**). Our *Psp*Cas13b design of potent crRNAs showed ∼87.8% average silencing efficiency (EGFP and TagBFP together, **Figure 2c, 2f**) and outperformed *Rfx*Cas13d design, further validating the accuracy of our prediction tool (**Figure 2c & 2f**).

To further investigate the enrichment of a G-rich motif at the 5’end of potent crRNAs and C bases at the 5’end of ineffective crRNAs, we hypothesized that altering these sequences in a *bona fide* spacer sequence may either worsen or improve their silencing efficiency. First, we selected 11 crRNAs that possess a GG sequence at 1^st^ and 2^nd^ positions of the spacer which we altered to CC by spacer mutagenesis. The data showed substantial compromise in the silencing efficiency of the majority of these crRNAs (**Extended Data Fig. 5a**). We also mutated 3, 2, or 1 G base(s) at the 5’end of the spacer to a C residue(s) and found that the substitution of 3 or 2 C bases at the 5’end of the spacer reduces their silencing by >99% and ∼70% respectively, while the introduction of a single C base at spacer position 1, 2, or 3 has no significant effect on the potency of the crRNA (**Extended Data Fig. 5b-5c**).

Next, we selected ineffective crRNAs lacking a GG sequence at their 5’end, and then modified them either by inserting an additional G at the first position, substituting the 1^st^ nucleotide to a G, or substituting the 1^st^ and 2^nd^ nucleotides to a GG (**Figure 2i-2o**). Importantly, the data demonstrated that G sequences at the 5’end of the spacer greatly increase the potency of crRNA despite the introduction of spacer-target mismatch (**Figure 2i-2o**). We questioned whether the improvement in silencing efficiency of crRNAs harbouring a G-rich motif at their 5’end could be secondary to changes in crRNA abundance. We quantified the expression levels of original crRNA or mutated crRNAs harbouring 5’end G motifs using quantitative real-time PCR (RT-PCR). Although not statistically significant, we observed an increase in crRNA abundance when a G-rich motif is present at the 5’end (**Extended Data Fig. 6**).

In addition to mCherry, we also show that C to G substitutions in key spacer positions (1, 2, 11, 15, 16, 17) can further improve the silencing efficiency of crRNAs targeting other transcripts (**Extended Data Fig. 7a-7l**). Indeed, when crRNA design choices are restricted, *de novo* design of crRNAs incorporating mismatched G bases at these key positions can substantially increase their potency despite introducing nucleotide mismatches with the target.

To facilitate the use of our optimized and validated spacer nucleotide-based formula for potent crRNA design, we created a user-friendly webpage (https://cas13b.github.io/) to assist the community with their silencing assays. This *in-silico* tool requires only the targeted sequence as input to create single-base tiled spacer sequences and rank them based on their predicted potency (see Methods).

### Comprehensive mutagenesis of spacer-target interaction

Understanding *Psp*Cas13b specificity, off-targeting potential, and its capability to discriminate between two transcripts that share extensive sequence homology is extremely important for evaluating the potential and the limitations of *Psp*Cas13-based RNA silencing. To study crRNA spacer promiscuity and the consequent *Psp*Cas13b targeting resolution, we conducted a comprehensive spacer mutagenesis to introduce mismatches with the target at various spacer positions. First, we introduced 3, 6, 9, 12, 15, 18, 21, 24, 27, and 30-nt successive mismatches starting from the 3’ and 5’ends of the spacer (**Figure 3a & 3b**). 3-nt mismatches at the 3’end of spacers (position 28-30) did not affect the silencing efficiency of this crRNA, whereas mismatches greater that 3-nt completely abrogated its silencing (**Figure 3a**). In contrast to the 3’end, all 5’end mismatches resulted in complete loss of silencing including 3-nt mismatches at the 5’ end (**Figure 3b**). Based on our earlier findings (**Extended Data Fig. 5**), silencing loss consequent to the introduction of a 3-nt mutation at the 5’end is likely attributable to the substitution of a GGG motif by a CCC sequence rather than spacer-target mismatch itself, thus reaffirming the importance of a G-rich motif at the 5’end of potent crRNAs as previously described (**Figure 1 & Figure 2**).

**Figure 3.**
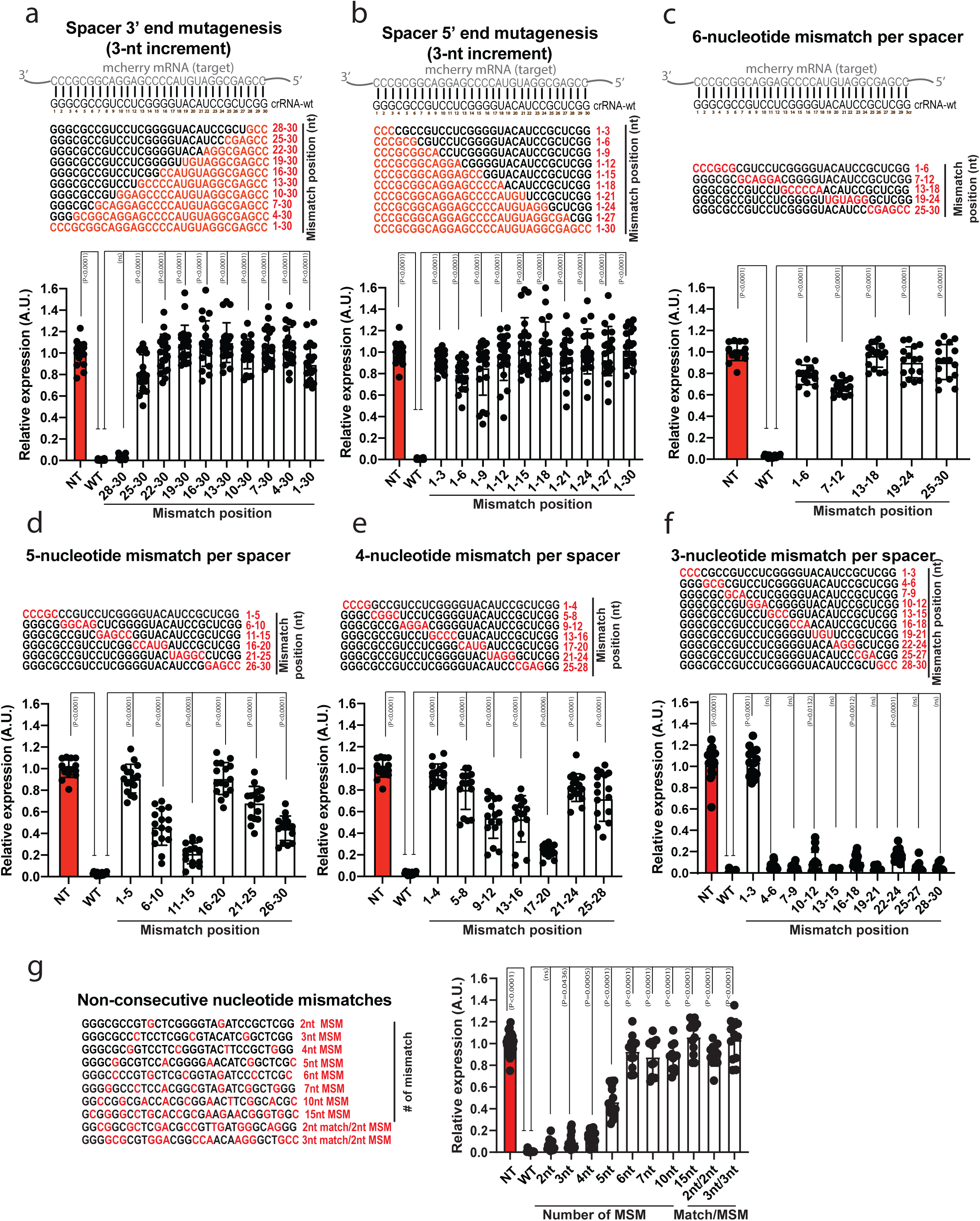
Comprehensive mutagenesis of *Psp*Cas13b spacer-target interaction revealed the interface between mismatch tolerance and loss of activity. **(a–g)** Comprehensive analysis of spacer-target interaction examining specificity and mismatch tolerance of *Psp*Cas13b at various positions of crRNA spacer sequence. Perturbation of spacer-target interaction through spacer mutagenesis to introduce 3, 6, 9, 12, 15, 18, 21, 24, 27, and 30-nucleotide consecutive mismatch at the **(a)** 3’ end and **(b)** 5’ end of the spacer. Perturbation of spacer-target interaction through spacer mutagenesis to introduce **(c)** 6, **(d)** 5, **(e)** 4, and **(f)** 3-nucleotide consecutive mismatch at various positions of the spacer. **(g)** Perturbation of spacer-target interaction through spacer mutagenesis to introduce various numbers of non-consecutive nucleotide mismatches at various positions. The nucleotides in red highlight mismatch positions in the spacer sequence. Data points in the graph are mean fluorescence from 4 representative field of views per condition imaged; *N* = 3 or 4. The data are represented in arbitrary units (A.U.). Errors are SD and *p*-values of one-way Anova test are indicated (95% confidence interval). *N* is the number of independent biological replicates. Source data are provided as a Source data file.

We also created crRNA constructs harbouring 6-nt, 5-nt, 4-nt, and 3-nt mismatches at different spacer positions and probed their silencing efficiency in live cells (**Figure 3c-3f**). Overall, 6-nt mismatches largely compromised the efficiency of *Psp*Cas13b regardless of mismatch position (**Figure 3c**). 5-nt mismatches at positions 6-10, 11-15, and 26-30 exhibited a partial loss of silencing, while mismatches at positions 1-5, 16-20, and 21-25 led to a near complete or complete loss of silencing (**Figure 3d**). 4-nt mismatches at positions 9-12, 13-16, and 17-20 retained partial silencing activity, whereas mismatches at positions 1-4, 5-8, 21-24, and 25-28 yielded a complete loss of silencing (**Figure 3e**). Notably, crRNA constructs harbouring 3-nt mismatches at various spacer positions were well tolerated and yielded no or minor loss of silencing, except for mutations at position 1-3 that, as anticipated, led to a total loss of silencing likely due to 5’end GGG removal (**Figure 3f**).

Whilst the preceding experiments established the tolerance for consecutive spacer-target mismatches, we questioned whether the silencing profile of non-consecutive mismatches may differ. We destabilized the spacer-target interaction by introducing 2, 3, 4, 5, 6, 7, 10, and 15 non-consecutive mismatches spread throughout the spacer (**Figure 3g**). We noticed that 2, 3, and 4 non-consecutive mismatches were tolerated and led to negligible loss of silencing. However, more than 4-nt non-consecutive mismatches led to a substantial or complete loss of silencing. Likewise, multiple successive 2 or 3 nucleotide mismatches spread throughout the spacer sequence also completely abolished its silencing activity (**Figure 3g**). These data revealed the targeting resolution of *Psp*Cas13b and suggest that >4-nt non-consecutive mismatches critically destabilise spacer-target interaction and compromise *Psp*Cas13b activity. In addition, the data also suggest that endogenous targets with partial sequence homology are unlikely to be impacted by off-target silencing due to the required minimum ∼25 consecutive or non-consecutive nucleotide basepairing. These mutagenesis data provide further evidence that highly effective crRNAs can be readily designed with minimal or no off-target effects.

### Efficient silencing of oncogenic fusion drivers

Now that we revealed key design principles of *Psp*Cas13b, we questioned whether we can reprogram this CRISPR enzyme to silence major oncogenic gene fusion transcripts. The breakpoint at the interface between the two genes offers a unique targetable sequence at the RNA level, which remains largely unexplored. We designed 21 tiled crRNAs (3-nucleotide resolution) targeting the breakpoint of 3 oncogenic gene fusions BCR-ABL1, SFPQ-ABL1, and SNX2-ABL1 that are established drivers of acute lymphoblastic leukemias (ALL)^24–26^. The gene fusions were each cloned into an IRES-GFP vector that produces the-gene-of-interest-IRES-GFP transcript, which is subsequently translated into separate proteins due to the presence of the IRES sequence. Therefore, an efficient targeting of the gene fusion transcript by *Psp*Cas13b is anticipated to lead to loss of GFP fluorescence due to sequence-specific recognition, cleavage, and degradation of the fusion-GFP transcript. Overall, microscopy data from 3-nucleotide resolution tiled crRNAs showed high silencing efficiency of all 3 gene fusions, although, once more the silencing efficiency varied depending on the position of the crRNA (**Figure 4a-4c**). Analysis of mRNA levels of gene transcripts by RT-qPCR confirmed high silencing efficiency with numerous crRNAs, although the magnitude of variance between crRNAs was less pronounced than suggested by the microscopy assay (**Figure 4d-4f**), possibly due to an additional Cas13-mediated protein translation regulation. Western blot analysis of the BCR-ABL1 protein expression further confirmed high silencing of BCR-ABL1 at the protein level, which, consistent with the microscopy data, was dependent on the position of crRNAs tested, with −12, −9 and +12 crRNAs exhibited the highest silencing efficiencies (**Figure 4g)**. Analysis of STAT5 and ERK phosphorylation, a hallmark of BCR-ABL1 dependent oncogenic signalling (**Figure 4h**), confirmed that potent crRNAs can efficiently suppress BCR-ABL1 and its downstream oncogenic networks (**Figure 4i**). Imatinib, a small inhibitory molecule that blocks the tyrosine kinase domain of ABL1 (**Figure 4h**), inhibited BCR-ABL1 mediated phosphorylation of STAT5 and ERK without altering the expression levels of BCR-ABL1 protein, whereas *Psp*Cas13b crRNAs efficiently silenced BCR-ABL1 protein expression and the downstream phosphorylation of STAT5 and ERK (**Figure 4i**). Interestingly, the most potent crRNA+12 showed greater suppression of STAT5 phosphorylation than Imatinib, consistent with its high efficacy in depleting the BCR-ABL1 protein through mRNA silencing (**Figure 4i**).

**Figure 4.**
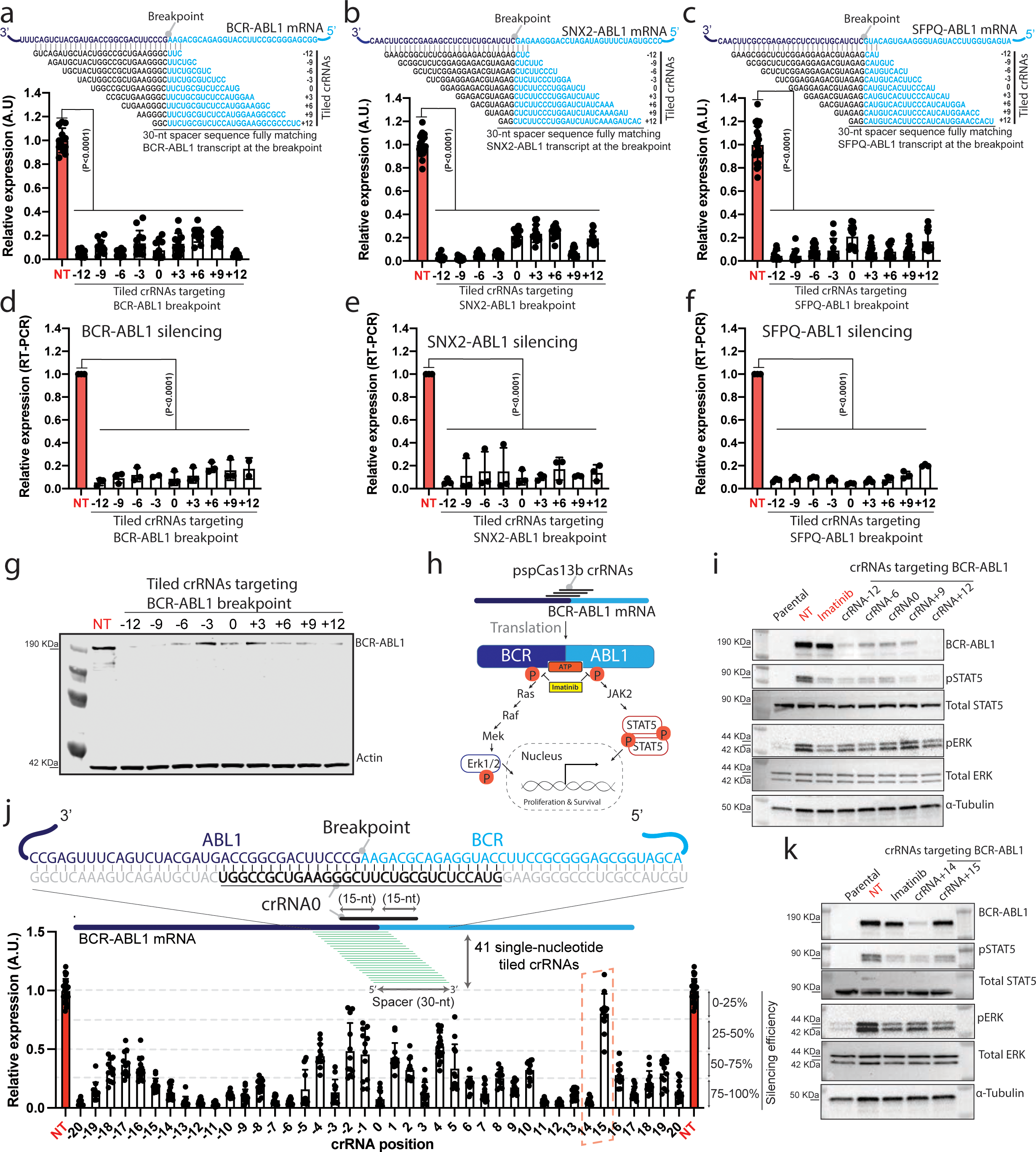
Reprogrammed *Psp*Cas13b suppresses fusion gene transcripts with high efficiency. **(a-c)** EGFP reporter assay to assess the silencing efficiency of tiled *Psp*Cas13b crRNAs with 3-nucleotide resolution targeting the breakpoint region of fusion transcripts BCR-ABL1 **(a**), SNX2-ABL1 **(b)** and SFPQ-ABL1 **(c)**. Data points in the graphs are mean fluorescence from 4 representative fields of view per condition imaged; *N* = 3. The data are represented in arbitrary units (A.U.). Errors are SD and *p*-values of one-way Anova test are indicated (95% confidence interval). **(d-f)** RT-PCR assays measuring silencing efficiency of tiled *Psp*Cas13b crRNAs (3-nucleotide resolution) targeting the breakpoint regions of fusion transcripts BCR-ABL1 **(d)**, SNX2-ABL1 **(e)** and SFPQ-ABL1 **(f)**; *N* = 3; Data are normalized means and errors are SEM; Results are analysed by one-way Anova test with *p*-values indicated (95% confidence interval). **(g)** Representative Western blot analysis to examine the expression level of BCR-ABL1 protein in HEK 293T cells expressing tiled crRNAs with 3-nucleotide increment targeting the breakpoint region of BCR-ABL1 transcripts 24 h post-transfection; *N* = 3. (See uncropped blots from 3 independent biological experiments in Source file). **(h)** Schematic of BCR-ABL1 dependent phosphorylation of ERK and Stat proteins, and inhibition of BCR-ABL1 oncogenic activity with Imatinib, a potent competitor of ATP binding to ABL kinase. **(i)** Representative Western blot analysis to examine the suppression of BCR-ABL1 expression and the subsequent inhibition of STAT5 and ERK phosphorylation in HEK 293T cells expressing BCR-ABL1, *Psp*Cas13b and either NT or crRNA targeting the BCR-ABL1 24 h post-transfection. HEK 293T cells expressing BCR-ABL1 and *Psp*Cas13b treated with 1µM imatinib for 4 hours were used as a positive control. Parental cells are HEK 293T cells transfected with *Psp*Cas13b, NT and a random control plasmid. This condition shows the baseline expression of pSTAT5 and pERK in BCR-ABL1 independent manner; *N* = 2. **(j)** 41 single-nucleotide tiled crRNAs targeting the mRNA region surrounding the breakpoint of BCR-ABL1 to reveal the relationship between RNA sequence, accessibility, and *Psp*Cas13b silencing efficiency. The schematic shows the sequence of BCR-ABL1 RNA covered by 41 tiled crRNAs and RNA-RNA duplex formed by spacer-target interaction. The red dashed box highlights two adjacent crRNAs (14 & 15) with markedly contrasted silencing efficiency. Data points in the graph are normalized mean fluorescence from 4 representative fields of view imaged in *N* = 3. The data are represented in arbitrary units (A.U.). Errors are SD with 95% confidence interval. **(k)** Representative western blot analysis to examine the silencing efficiency of single-base resolved crRNAs 14 & 15 that target BCR-ABL1 mRNA. crRNA potency is examined through the silencing of BCR-ABL1 protein and phosphorylation of STAT5 and ERK proteins. Cells expressing BCR-ABL1, *Psp*Cas13b and either NT or crRNA targeting the BCR-ABL1 were harvested for WB analysis 24 h post-transfection. 1µM imatinib treatment for 4 hours was used as a positive control to inhibit BCR-ABL1 kinase activity. Parental cells are HEK 293T cells transfected with *Psp*Cas13b, NT and a control plasmid to examine the baseline expression of pSTAT5 and pERK in a BCR-ABL1 independent manner; *N* = 3. (See uncropped blots from 3 independent biological experiments in Source file). *N* is the number of independent biological experiments. Source data are provided as a Source data file.

Next, we sought to investigate whether single-nucleotide tiled crRNAs targeting BCR-ABL1 would also show highly variable levels of silencing. We cloned and deployed 41 individual tiled crRNAs across the breakpoint of BCR-ABL1 (**Figure 4j**). Again, we observed that the silencing efficiency highly varied even between neighbouring crRNAs. For instance, despite 96.6% sequence homology and only a single nucleotide position shift, crRNA14 achieved >90% silencing while crRNA15 exhibited no silencing, with consistent results evident in both quantitative microscopy and Western blot analyses (**Figure 4j & 4k**). The potent crRNA+14 also exhibited higher silencing of downstream STAT5 phosphorylation (**Figure 4k**). The contrasted silencing activity obtained with single-base resolved crRNAs within the same targeted region confirms the presence of key RNA sequences or features that profoundly influence *Psp*Cas13b activity.

Taken together, these data demonstrated the utility of *Psp*Cas13b as a versatile tool to efficiently silence tumour drivers such as fusion transcripts and alter their oncogenic signalling networks, and highlight the importance of rational design for maximum crRNA potency.

### Absolute discrimination between fusion and wild type RNAs

Previous spacer mutagenesis experiments indicated that *Psp*Cas13b can discriminate between two RNAs that share extensive sequence homology (**Figure 3**). We questioned whether this discriminatory resolution is generalisable to cancer-specific oncogenic RNAs. To determine this, we introduced 3, 4, 5, 6, 7, 10, and 14 non-consecutive mismatches between the spacer of BCR-ABL1 crRNA (crBCR-ABL1) and the targeted breakpoint sequence (**Figure 5a**). The data revealed that 3 nucleotide mismatches were well tolerated. However, 4 or higher number of non-consecutive nucleotide mismatches drastically impaired crRNA silencing efficiency (**Figure 5a**). 3 consecutive nucleotide mismatches at various positions did not affect the silencing of BCR-ABL1. 6 consecutive nucleotide mismatches at the 3’end (25-30) or at the central region (12-17) led to notable loss of silencing, while 9 consecutive nucleotide mismatches dramatically curtailed silencing irrespective of position (**Figure 5b**). Western blot analysis of BCR-ABL1 protein expression confirmed these data and showed that 3-nucleotide mismatches are well tolerated, while 4-nucleotide mismatches or higher led to substantial or complete loss of silencing (**Figure 5c**). Overall, the data highlights the specificity of *Psp*Cas13b and its potential to discriminate between transcripts despite extensive sequence homology.

**Figure 5.**
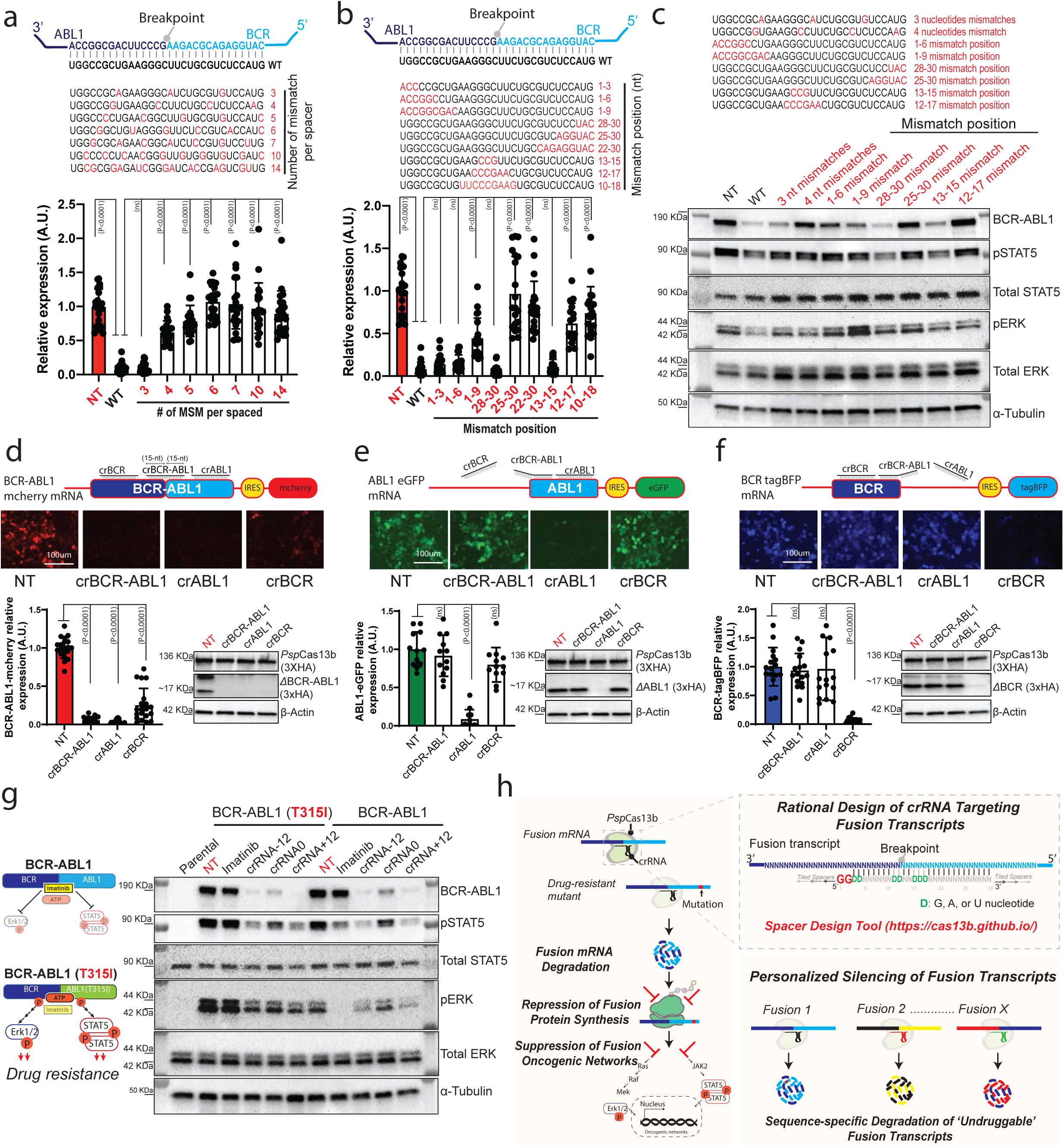
The targeting of the breakpoint can efficiently discriminate between translocated tumour RNAs and wild type variants despite extensive sequence homology. **(a–b)** Comprehensive analysis of spacer-target interaction examining specificity and mismatch tolerance of *Psp*Cas13b crRNAs targeting the breakpoint region of BCR-ABL1 transcript. The nucleotides in red highlight various mismatch positions in the spacer sequence. Data points in the graph are mean fluorescence from 4 representative field of views per condition imaged; *N* = 4. The data are represented in arbitrary units (A.U.). Errors are SD and *p*-values of one-way Anova test are indicated (95% confidence interval). **(c)** Western blot analysis examining the expression level of BCR-ABL1 protein and phosphorylation status of STAT5 and ERK in HEK 293T cells expressing crRNAs with various mismatches 24 h post-transfection; *N* = 3. (See uncropped blots from 3 independent biological experiments in Source file). **(d-f)** 3 colour fluorescence-based reporter assays to assess the on-target specificity of crRNA targeting the breakpoint region of BCR-ABL1 and potential off-targeting of wild type BCR and ABL1 transcripts in HEK 293T cells 48 h post-transfection. The schematics show BCR-ABL1-mCherry mRNA **(d)**, ABL1-eGFP mRNA, **(e)** and BCR-TagBFP mRNA **(f)** and their interaction with crBCR, crBCR-ABL1 and crABL1crRNAs through full, partial, or no spacer-target basepairing. Representative fluorescence microscopy images show the silencing efficiency of crBCR, crBCR-ABL1 and crABL1 crRNAs targeting BCR-ABL1-mCherry **(d)**, ABL1-eGFP **(e)** and BCR-TagBFP transcripts **(f)** in HEK 293T cells. Scale bar = 100 μm. Similar results were obtained in 3 independent experiments. Unprocessed representative images are provided in the Source Data file. The histograms quantify the silencing of BCR-ABL1-mCherry **(d)**, ABL1-eGFP **(e)** and BCR-TagBFP **(f)**. Data points are normalized mean fluorescence from 4 representative fields of view per condition imaged. The data are represented in arbitrary units (A.U.). Errors are SD and *p*-values of one-way Anova test are indicated (95% confidence interval). *N* = 3. The right panels show representative western blot analyses to examine the expression levels of ΔBCR-ABL, ΔABL1, and ΔBCR proteins from data in **d-e;** *N* = 3. **(g)** Representative western blot analysis to examine the suppression of *Imatinib*-resistant T315I BCR-ABL1 with *Psp*Cas13b. The schematic (left) illustrates *Imatinib*-sensitivity or *Imatinib*-resistance of WT and T315I variants, respectively. BCR-ABL1 expression and STAT5/ERK phosphorylation status were examined in HEK 293T cells expressing WT or T315I BCR-ABL1 variants, *Psp*Cas13b and either NT or crRNAs targeting the BCR-ABL1 breakpoint 24 h post-transfection. HEK 293T cells expressing BCR-ABL1 variants and *Psp*Cas13b were treated with 1µM imatinib for 4 hours as a positive control. Parental cells are HEK 293T cells transfected with *Psp*Cas13b, NT and a control plasmid, which shows the baseline expression of pSTAT5 and pERK in BCR-ABL1 independent manner; *N* = 3. *N* is the number of independent biological replicates. Source data are provided as a Source data file. **(h)** Schematic summarizing the main findings of this study. The left panel shows that rational design of crRNAs targeting the breakpoint of gene fusion transcripts enables efficient and specific silencing of these oncogenic drivers as well as their related drug-resistant mutants. *Psp*Cas13b-mediated RNA degradation suppresses fusion protein translation and their downstream oncogenic networks. The top right panel summarizes spacer nucleotide positions that are important for rational design of potent crRNA, which are integrated into our *in-silico* tool (https://cas13b.github.io/). The bottom right panel shows the versality and therapeutic potential of *Psp*Cas13b to silence various ‘*undruggable*’ gene fusion transcripts and their oncogenic networks in a personalised manner.

To confirm this specificity, we tested crRNAs targeting BCR-ABL1 fusion against wild type non-translocated BCR and ABL1 transcripts which are expressed in normal tissues. We cloned constructs encoding partial mRNA sequences of the BCR-ABL1 fusion, ABL1 alone, and BCR alone in frame with mCherry, eGFP, or TagBFP fluorescent reporters, respectively (**Figure 5d-5f**). We designed 3 crRNAs targeting the BCR-ABL1 breakpoint sequence (crBCR-ABL1), BCR sequence (crBCR), or ABL1 sequence (crABL1) that we tested against the aforementioned constructs. The fluorescence signals from mCherry, eGFP, and TagBFP enable accurate quantification of on-target and off-target silencing with these crRNAs. As anticipated, all 3 crRNAs silenced the *bona fide* BCR-ABL1 transcript as this mRNA possesses full-length spacer binding sites for all three crRNAs (**Figure 5d**). However, ABL1 and BCR transcripts were silenced only by their cognate crABL1 and crBCR crRNAs (**Figure 5e-5f**). Notably, crBCR-ABL1 targeting the breakpoint sequence had no effect on either BCR or ABL1 wildtype transcripts despite 15-nucleotide sequence basepairing (**Figure 5e-5f**). Western blot analysis confirmed these data (**Figure 5d-5f**), demonstrating the high-resolution capability of *Psp*Cas13b and its utility to specifically silence oncogenic gene fusion drivers at the RNA level while sparing non-translocated wild type transcripts.

### High potency against drug-resistant point-mutated fusion transcripts

Acquired drug resistance to all approved ABL1 kinase inhibitors through secondary mutations remains a major challenge in the treatment of BCR-ABL1 driven leukemias^27^. For instance, the BCR-ABL1 kinase domain mutation Thr315Ile (T315I) confers resistance to *Imatinib* and drives tumour relapse^28^. We hypothesised that unlike *Imatinib*, targeting the breakpoint of BCR-ABL1 transcript with potent crRNAs will remain effective against both BCR-ABL1 variants as the mutation is located outside the targeted sequences at the breakpoint. We tested the potency of either the *Imatinib* or three *Psp*cas13b crRNAs targeting the breakpoint of BCR-ABL1. As anticipated, *Imatinib* efficiently inhibited the oncogenic signalling of ancestral BCR-ABL1 but failed to effectively suppress T315I BCR-ABL1 downstream signalling (**Figure 5g**). Notably, all three *Psp*Cas13b crRNAs we tested largely inhibited the expression of ancestral and T315I BCR-ABL1 proteins and their downstream oncogenic signalling as exemplified by phospho-STAT5 and phospho-ERK inhibition. Consistent with previous data, crRNA-12 and crRNA+12 achieved the highest inhibitory effect due to higher silencing potency (**Figure 5g**). These data demonstrate that targeting the breakpoint of BCR-ABL1 transcript can overcome drug resistance commonly observed in recurrent leukemia.

## DISCUSSION

Our study elucidates key molecular principles of *Psp*Cas13b RNA silencing that enabled *de novo* design of crRNAs for a potent suppression of fusion oncogenic transcripts, including variants that drive clinical drug resistance.

Overall, Cas13 nucleoproteins can offer effective and specific silencing of targeted transcripts without the risk of permanent alteration of genomic DNA^19–21^. Our single-nucleotide resolution tiled crRNA screens and the comprehensive *in silico* analysis demonstrated that, unlike other CRISPR systems^29^, *Psp*Cas13b silencing is not restricted by a defined PFS (PAM-like) sequence, underlining its greater design flexibility (**Figure 1**). Among the strongest predictors of Cas13 activity, we identified a ‘G-rich’ motif at the 5’end of the spacer as key determinant of crRNA potency (**Figure 1**). Notably, *de novo* designed crRNAs harbouring target matched or target-mismatched ‘GG’ sequence at the 1^st^ and 2^nd^ nucleotide positions of the spacer can greatly enhance the silencing potency of otherwise poorly effective crRNAs (**Figure 2**). The ability of this target mismatched ‘GG’ motif to rescue the potency of certain ineffective crRNAs expands the range of effective crRNAs for a given target, which is particularly important for narrowly defined target sequences, especially when targeting breakpoint region of fusion transcripts (**Figure 4&5**), RNA isoforms, or single-nucleotide variants. We also revealed ‘C’ nucleotides at the 5’end or central spacer positions that can reduce the efficacy of crRNAs (**Figure 1&2; Extended Data Fig. 5**). The substitution of these ‘C’ bases with ‘G’ can enhance crRNA potency despite the introduction of mismatches that do not exceed 3 (**Figure 2; Extended Data Fig. 7**). Integrating these discoveries into our design process enabled us to consistently generate highly potent crRNAs with an average silencing efficiency of ∼87% (**Figure 2**), outperforming the benchmark prediction algorithm available for *Rfx*Cas13d ortholog^30^. Therefore, we created an open access and user-friendly webpage (https://cas13b.github.io/) to assist a wider community with their design of predicted potent crRNAs for *Psp*Cas13b using the aforementioned criteria.

The Specificity of spacer-target interaction is core to the capability of *Psp*Cas13b to selectively silence target RNAs in the RNA-crowded cellular environment^31^, especially in the context of sequence homology between various RNAs (**Figure 3**). Although some *Psp*Cas13b spacer sequences can tolerate ∼3 to 5 consecutive or non-consecutive nucleotide mismatches with the target, the 30-nucleotide spacer length enables this protein to retain remarkably high specificity (**Figure 3**). Thus, this study revealed the interface between mismatch tolerance and intolerance, and provides a blueprint for silencing various pathogenic transcripts with single-base resolution, further expanding the targeting spectrum to point-mutated variants^32^.

The most important aspect of *Psp*Cas13b-mediated targeting is its translational potential to silence any treatment-resistant or ‘*undruggable*’ oncogenic gene fusions^1, 7, 32^. Over 3,297 unique fusion transcripts across multiple cancer types have been described^32^, most of which remain ‘*undruggable*’ but could be targeted by personalized *Psp*Cas13b drugs. Indeed, we show that *Psp*Cas13b can efficiently recognise and silence three different fusion transcripts including BCR-ABL1, a well-established driver of leukemias^25, 33–36^, without the off-targeting of non-translocated wildtype variants that are expressed in non-malignant tissues (**Figure 4 & Figure 5**). BCR-ABL1 transcript silencing led to subsequent depletion of the fusion protein and thereby inhibited the phosphorylation and activation of downstream STAT5 and ERK signalling pathways that are a hallmark of BCR-ABL1 driven cancers, which outperformed the efficiency of the tyrosine kinase inhibitor *Imatinib* (**Figure 4**). The use of *imatinib* in treating BCR-ABL1-positive leukemias can be associated with severe toxicity due to the non-specific inhibition of ABL1 and ABL1-like kinases in normal tissues^37^. Conversely, *Psp*Cas13b-mediated targeting of the fusion breakpoint shows superior specificity without the off-targeting of non-translocated BCR, ABL1, or other ABL1-like kinases (**Figure 4 & 5**), which is anticipated to exhibit very mild or reduced toxicity *in vivo* compared to *Imatinib*.

In addition to drug toxicity, acquired resistance through secondary mutations remains a major challenge in the management of cancers. For example, the BCR-ABL1 kinase domain “gatekeeper” mutation T315I confers resistance to first-line ABL1 inhibitors, including *Imatinib,* and often drives tumour relapse in patients^28^. Unlike *Imatinib*, potent *Psp*Cas13b crRNA targeting the breakpoint of BCR-ABL1 transcript remained highly effective against both wildtype and T315I BCR-ABL1 variants, and suppressed their downstream oncogenic networks (**Figure 5g**). These data demonstrate the great potential of this *Psp*Cas13b tool to silence ‘*undruggable*’ fusion transcripts that drive various tumours and overcome their mutation-driven drug resistance (**Figure 5h**).

Excitingly, the breakpoint sequence of a given fusion driver can be easily obtained from patients’ tumour RNA sequencing^32^, which would enable the manufacturing of personalized *Psp*Cas13b drugs in just few days (**Figure 5h**), making these drug development pipelines suitable for cancer discovery as well as potential clinical translation to suppress various ‘*undruggable*’ tumour drivers in actionable time frame.

As a conclusion, *Psp*Cas13b design flexibility, specificity, and the apparent lack of collateral activity (**Extended Data Fig. 7m**) provide a new conceptual framework that could transform the landscape of personalized medicine through targeting ‘*undruggable’* or drug-resistant pathogenic RNAs that drive human disorders including cancers (**Figure 5h**).

## Acknowledgment

The authors thank all lab members from the Trapani, Ekert, Voskoboinik, and Fareh labs for facilitating experiments and discussions. This work was supported by a Cancer Council Victoria Ventures grant (ID 829606) to M.F, P.G.E, and J.A.T., by a Peter MacCallum Cancer centre strategic plan funding in partnership with the Children’s Cancer Institute Australia (CCIA) to M.F, by a Peter MacCallum Foundation grant to M.F (ID 2119), and by a the National Health and Medical Research Council (NHMRC) of Australia through a program grant to J.A.T.

## Authors’ contribution

M.F conceived the study. M.F, J.A.T, and P.G.E supervised the study. W.H and M.F designed the experiments. W.H, and M.F, performed the experiments and analysed the data. T.S. cloned Imatinib-resistant BCR-ABL1 gene fusion construct and helped W.H with WB experiments. A.K, S.Q, and W.H performed the computational analysis with input from M.F. A.K and D.M created the crRNA design code and webpage with input from W.H and M.F. All authors discussed the project and the data. W.H and M.F generated graphs, figures and wrote the manuscript. W.H, J.A.T, P.G.E, J.M.L.C, M.H, and M.F revised and edited the manuscript. All authors read, commented, edited, and approved the manuscript.

### Data availability

All data are available in the main text and supplementary materials. Source Data are provided with this paper. All key plasmids constructed in this study, their sequences, and maps will be deposited to Addgene upon publication.

## Code Availability

The bioinformatic codes for the design of predicted potent crRNA is available at https://github.com/david-ma/cas13. crRNA design web-server described here is available at: https://cas13b.github.io/.

## Competing interests

The other authors declare no conflict of interest.

## METHODS

### Design and cloning of crRNAs for *Psp*Cas13b

The design and cloning of *Psp*Cas13b crRNAs were according to a previous publication^14^. Briefly, individual guide RNAs were cloned into the pC0043-*Psp*Cas13b crRNA backbone (addgene#103854, a gift from Feng Zhang lab, later referred to as crRNA backbone) which contains a *Psp*Cas13b crRNA direct repeat (DR) sequence and two BbsI restriction sites for the cloning of spacer sequence. A total of 20 μg crRNA backbone was digested by BbsI restriction enzymes (NEB, R3539) following the manufacturer’s instructions for 2 hours at 37C°. Backbone linearization was checked by 1% agarose gel. The digested backbone was purified with NucleoSpin Gel and PCR Clean-up Kit (Macherey-Nagel, 740609.50), aliquoted, and stored in −20C° until using.

For crRNA cloning, forward and reverse single-stranded DNA oligonucleotides containing CACC and CAAC overhangs respectively, were ordered from Sigma or IDT (100 μM). A total of 1.5 μL of 100 μM the forward and reverse DNA oligos were annealed in 47 μL annealing buffer (5 μl NEB buffer 3.1 and 42 μL H_2_O) by 5 min incubation at 95 °C and slow cool down in the heating block overnight. 1 μL of the annealed oligos were ligated with 0.04 ng digested *Psp*CAs13b crRNA backbone in 10 μL of T4 ligation buffer (3 h, RT) (Promega, M1801). All *Psp*Cas13b crRNA spacer sequences used in this study are listed in Supplementary File 1. All crRNAs and *Psp*Cas13b clones that are generated in this study were verified by Sanger sequencing (AGRF, AUSTRALIA). The primers used for PCR and Sanger sequencing are listed in Supplementary File 3.

### Cloning of BCR-ABL1 T315I, BCR-ABL1, ABL1 and BCR fragments

The partial sequences of BCR-ABL1, ABL1 and BCR were designed according to the full length BCR-ABL1 P190. The IDT DNA synthesis platform provided the three sequences that were subsequently cloned into MSCV-IRES-mCherry, MSCV-IRES-eGFP and MSCV-IRES-tagBFP vectors respectively in frame with 3xHA tag using EcoRI/BamHI digestion (Promega, R6011/Promega, R6021), gel purification, and ligation with T4 DNA ligase. The ligated product was transformed into chemically competent bacteria (TOP10 or Stbl3) and positive clones were screened by PCR and Sanger sequencing (AGRF, AUSTRALIA). The BCR-ABL1-3xHA-IRES-mCherry, BCR-3xHA-IRES-tagBFP and ABL1-3xHA-IRES-EGFP constructs are available through Addgene (upon publication). The sequences of these DNA constructs are available in Supplementary File 2. The primers used for PCR and Sanger sequencing are listed in Supplementary File 3.

To generate the BCR-ABL1 T315I mutant construct, site-directed mutagenesis was performed by using the Phusion Site-Directed Mutagenesis Kit (Thermo Fisher F541) with the following primers with 5’ phosphorylation: (T315I_F: 5’CCCCGTTCTATATCATCATCGAGTTCATGACCTAC3’ and T315I_R: 5’GCTCCCGGGTGCAGACCCCAAGGAG 3’).

### Plasmid amplification and purification

The plasmid amplification and purification are described in a previous publication^16^. Briefly, TOP10 or Stbl3 bacteria were used for transformation. A total of 5–10 μL ligated plasmids were transformed into 30 μL of chemically competent bacteria by heat shock at 42 °C for 45 s, followed by 2 min on ice. The transformed bacteria were incubated in 500 μL LB broth media containing 75 μg/mL ampicillin (Sigma-Aldrich, A9393) for 1 h at 37 °C in a shaking incubator (200 rpm). The bacteria were pelleted by centrifugation at 6,000 rpm for 1 min at room temperature (RT), re-suspended in 100 μL of LB broth, and plated onto a pre-warmed 10 cm LB agar plate containing 75 μg/mL ampicillin, and incubated at 37 °C overnight. Next day, single colonies were picked and transferred into bacterial starter cultures and incubated for ∼6 h for mini-prep (Macherey-Nagel, NucleoSpin Plasimd Mini kit for plasmid DNA, 740588.50) or maxi-prep (Macherey-Nagel, NucleoBond Xtra Maxi Plus, 740416.50) DNA purification according to the standard manufacturer’s protocol.

### Cell culture

HEK 293 T (ATCC CRL-3216) cell line were cultured in DMEM high glucose media (Thermo Fisher, 11965092) containing 10% heat-inactivated fetal bovine serum (Thermo Fisher, 10100147), 100mg/ml Penicillin/-Streptomycin (Thermo Fisher, 151401220), and 2mM GlutaMAX (Thermo Fisher, A1286001). HEK 293 T cells were maintained at confluency between 20 and 80% in 37 °C incubators with 10% CO_2_. Cells were routinely tested and were mycoplasma negative.

### RNA silencing assays

All transfection experiments were performed using an optimized Lipofectamine 3000 transfection protocol (Thermo Fisher, L3000015). For RNA silencing in HEK 293 T, cells were plated at approximately 30,000 cells/100 μL/96-well in tissue culture treated flat-bottom 96-well plates (Corning) 18 h prior to transfection. For each well, a total of 100 ng DNA plasmids (22 ng of Ef1a-*Psp*Cas13b-NES-3xFLAG-T2A-BFP (addgene #173029) or pC0046-EF1a-*Psp*Cas13b-NES-HIV (addgene #103862), 22 ng crRNA plasmid, and 56 ng of the target gene) were mixed with 0.2 μL P3000 reagent in Opti-MEM Serum-free Medium (Thermo Fisher, 31985070) to a total of 5 μL (mix1). Separately, 4.7 μL of Opti-MEM was mixed with 0.3 μL Lipofectamine 3000 (mix2). Mix1 and mix2 were added together and incubated for 20 min at room temperature, then 10 μL of transfection mixture was added to each well. Supplementary File 4 summarizes the transfection conditions used in 96, 24, and 12-well plates. After transfection, cells were incubated at 37C°, 10% CO_2,_ and the transfection efficacy was monitored 24-72 hours post-transfection by fluorescent microscopy.

### Fluorescent microscopy analysis

For RNA silencing experiments, the fluorescence intensity was monitored using EVOS M5000 FL Cell Imaging System (Thermo Fisher). Images were taken 48 h post-transfection, and the fluorescence intensity of each image was quantified using a lab-written macro in ImageJ software. Briefly, all images obtained from a single experiment are simultaneously processed using a batch mode macro. First, images were converted to 8-bit, threshold adjusted, converted to black and white using Convert to Mask function, and fluorescence intensity per pixel measured using Analyze Particles function. Each single mean fluorescence intensity was obtained from four different field of views for each crRNA, and subsequently normalized to the non-targeting (NT) control crRNA. Two-fold or higher reduction in fluorescence intensity is considered as biologically relevant.

### Western Blot

Cells were washed three times with ice-cold PBS ± and lysed on ice in RIPA lysis buffer [50 mM Tris (Sigma-Aldrich, T1530), pH 8.0, 150 mM NaCl, 1% NP-40 (Sigma-Aldrich, I18896), 0.1% SDS, 0.5% sodium deoxycholate (Sigma-Aldrich, D6750)] containing protease inhibitor cocktail (Roche, 04693159001) and phosphatase inhibitor cocktail (Roche, 4906845001). Samples were incubated for 30min at 4 °C with rotation (25 rpm), and centrifuged at 16,000 g for 10 min, 4 °C. Supernatant was transferred to a new tube. Protein concentrations were quantified using the Pierce BCA Protein Assay Kit (Thermo Fisher, 23225) according to the manufacturer’s instructions. A total of 10 μg of protein diluted in 1x Bolt LDS sample buffer (Thermo Fisher, B007) and 1x Bolt sample reducing agent (Thermo Fisher, B009) were denatured at 95 °C for 5 min. Samples were resolved by Bolt Bis-Tris Plus 4–12% gels (Thermo Fisher, NW04120BOX) in 1x MES SDS running buffer (Thermo Fisher, B0002) and transferred to 0.45 μM PVDF membranes (Thermo Fisher, 88518) by a Trans-Blot Semi-Dry electrophoretic transfer cell (Bio-Rad) at 20 Volt for 30 min. Alternatively, samples were resolved by 4-15% Criterion TGX Precast Midi Protein gels (Bio-Rad, 5671084) in 1x Tris/glycine/SDS running buffer (Bio-Rad, 1610732) and transferred to 0.20 μM nitrocellulose membranes (Bio-Rad, 1704159) by a Trans-Blot Turbo Transfer System (Bio-Rad) with a HIGH MW protocol. Membranes were incubated in blocking buffer 5% (w/v) BSA (Sigma-Aldrich, A3059) in TBST with 0.15% Tween 20 (Sigma-Aldrich, P1379) for 1h at RT and probed overnight with primary antibodies at 4 °C. Blots were washed three times in TBST with 0.15% Tween20, followed by incubation with fluorophore-conjugated or HRP-conjugated secondary antibodies for 1h at RT. Membranes were washed in TBST (0.15% Tween20) three times and fluorescence or chemiluminescence was detected using the Odyssey CLx Imager 9140 (Li-cor), iBright CL1500 Imaging System (Thermo Fisher), or ChemiDoc Imaging System (Bio-Rad) . The antibodies used for western blots are listed in the Key Resources Table.

### RNA extraction, cDNA synthesis, and RT-PCR

Total RNA was isolated from around 5×10^5^ to 1×10^6^ cells using the NucleoSpin RNA Plus (MACHEREY-NAGEL, 740984.50) or Quick-RNA Miniprep Kit (Zymo Research, R1055) following the manufacturer’s instructions. For crRNA RT-PCR, total RNA was isolated by standard Trizol-chloroform extraction according to manufacturer’s instruction (Thermo Fisher, 15596026), followed by Dnase treatment with RQ1 RNase-Free DNase according to manufacturer’s instruction (Promega, M6101).

1μg total RNA was converted to cDNA using the high-capacity cDNA reverse transcription kit (Thermo Fisher, 4368814) following the manufacturer’s instructions. Quantitative RT-PCR reaction was performed in duplicate in a StepOne Real-Time PCR system (Thermo Fisher) using PowerUp™ SYBR™ Green Master Mix (Thermo Fisher, A25742). Total reaction mixture contains 0.2μl cDNA, 0.6μM forward primer and 0.6 μM reverse primer. Primers for RT-PCR are detailed in Supplementary File 3.

### Prediction of RNA secondary structure, RNA MFE, and RNA-RNA hybridization energy

RNAfold was used to predict the MFE of crRNA spacer, crRNA (DR and spacer), and the 70nt target region in the target RNA (20nt up/downstream from the 30nt-spacer-binding region). RNAfold was also used to explore the secondary structure of crRNAs and the target regions in the target RNAs. RNAplex^38^ and intaRNA^39^ were used to predict the hybridization energy and interaction energy between crRNA spacer and target RNA, respectively.

### *Psp*Cas13b crRNA design tool

The design of *in silico* prediction code is based on the experimental data presented in this article. Briefly, the algorithm generates single-nucleotide tiled spacer sequences for any input DNA or RNA sequence using HTML5, CSS and JavaScript libraries. The program then removes all spacer sequences that possess more than three consecutive T bases (>3T) that are predicted to act as a transcription termination signal and yield premature crRNAs. The algorithm scores the remaining spacer sequences based on their nucleotide composition and position. Spacers with a G nucleotide at the first or second positions receive a maximum score of +20 each. In contrast, a C nucleotide at spacer position 1, 2, 3, or 4 receive a penalty score of −20 each. Additionally, C bases at positions 11, 12, 15, 16 and 17 receive a −5 score each. All other nucleotides or spacer positions that did not show any distribution bias in the potent and ineffective crRNA receive a score of 0. The algorithm then calculates the cumulative score for each spacer and ranks them accordingly. As a result, the top spacers with high scores are enriched with G bases at 1^st^ and 2^nd^ positions, and depleted from C bases at positions 1, 2, 3, 4, 11, 12, 15, 16, and 17, and are predicted to yield potent silencing. Conversely, the lowest scoring spacers at the bottom of the list are enriched with C bases at positions 1, 2, 3, 4, 11, 12, 15, 16, and 17 and are predicted to yield ineffective silencing. The prediction accuracy of the algorithm is supported by in silico analysis and functional validation data in Figure 1**, 2, and Extended Data** Figure 2**, 3, 4, & 7**. This *Psp*Cas13b crRNA design tool is open source and available to the wider scientific community at https://cas13b.github.io/.

### Data analysis

Data analyses and visualizations (graphs) were performed in GraphPad Prism software version 9, unless stated otherwise. Specific statistical tests and numbers of independent biological replicates are mentioned in respective figure legends. The silencing efficiency of various crRNAs was analyzed using one-way Anova followed by Dunnett’s multiple comparison test where we compare every mean to a control mean as indicated in the Figures (95% confidence interval). The *P* values (P) are indicated in the Figures. *P* < 0.05 is considered statistically significant. Pearson correlation coefficient was used to analyze correlation between the crRNA silencing efficiency and potential parameters including crRNA MFE, target MFE, crRNA spacer MFE, crRNA-target RNA hybridization/interaction energy, crRNA spacer GC content, and A/U/G/C content. The R package ‘ggseqlogo’ was used to assess nucleotide preference in crRNA spacer and PFS sequences^40^. Delta probability graphs of spacer nucleotides were generated with Matplotlib^41^.

---

**Extended Data Figure 1.**
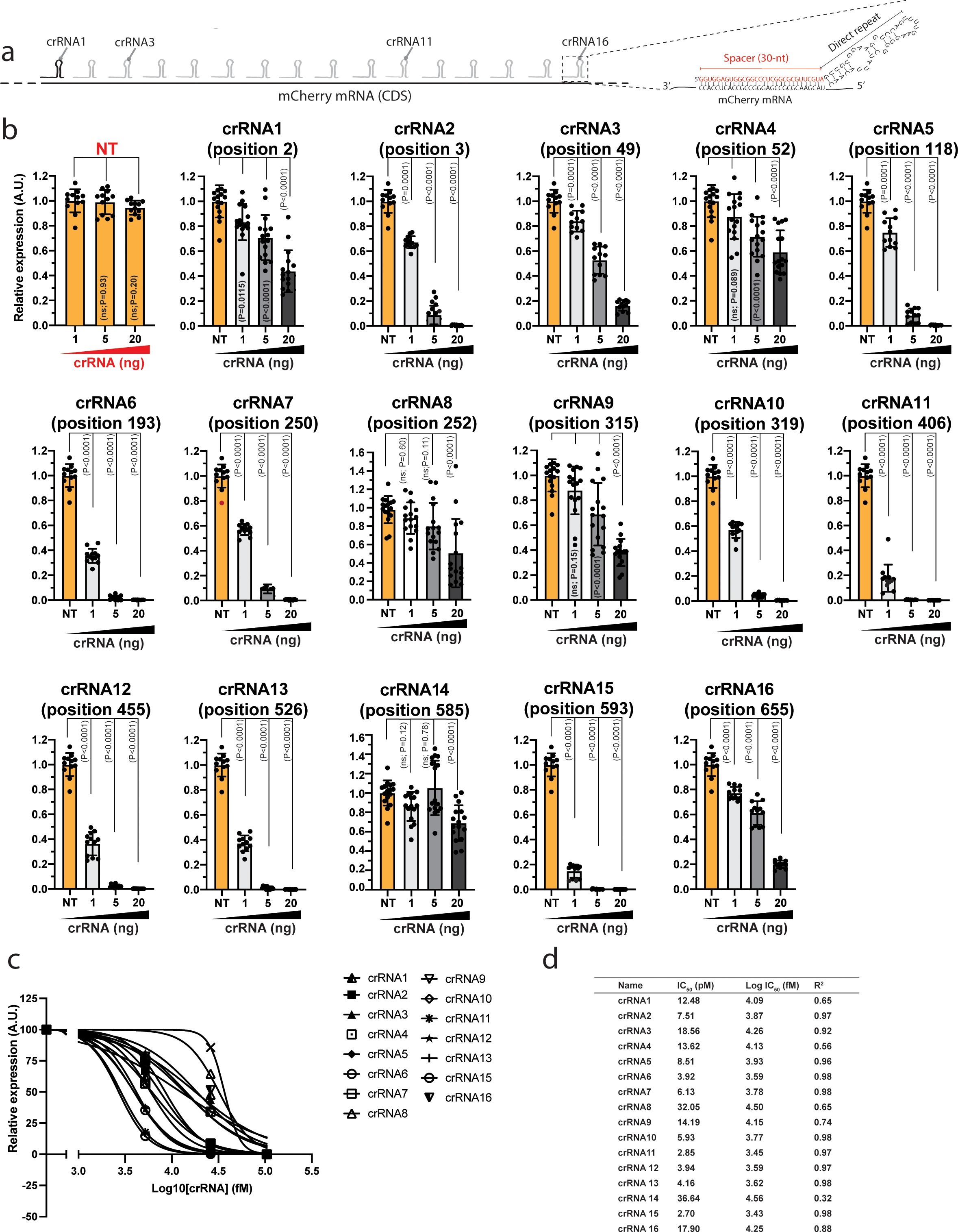
**(a)** Schematic of the 16 *Psp*Cas13b crRNAs targeting mCherry RNA. The detailed spacer and direct repeat of a representative crRNA is indicated in the right schematic. **(b)** crRNA dose-dependent silencing of mCherry transcripts with either NT (orange) or targeting crRNAs (grey); Data points in the graph are normalized mean fluorescence from 4 representative fields of view imaged in *N* = 3 or 4. The data are represented in arbitrary units (A.U.). Errors are SD and p-values of one-way Anova test are indicated (95% confidence interval). *N* is the number of independent biological experiments. Source data are provided as a Source data file. **(c)** Dose-response fitting of mCherry silencing data with non-targeting crRNA (NT) and 16 targeting crRNAs in **b**. **(d)** The IC_50_ values for 16 crRNAs targeting mCherry transcripts obtained from fitting in **c**. **Note:** We noticed marked differences in the silencing efficacy of the various crRNAs, even when they are designed to target neighbouring RNA locations. For example, crRNA6, crRNA11, crRNA12, crRNA13, and crRNA15 were extremely potent and degraded the majority of mCherry mRNA at a very low dose of 1 ng plasmid (5.2 pM). Conversely, crRNA1, crRNA4, crRNA8, crRNA9, and crRNA14 were inefficient and failed to completely degrade mCherry mRNA even at higher doses of 5 and 20 ng (26 and 104 pM) (**Extended Data Figure 1b**). crRNA potency was determined via calculation of the IC50 value, a dose that achieved 50% degradation of the target RNA, which confirmed the high variability in the silencing efficiency of various crRNAs (**Extended Data Figure 1a-1b**). Surprisingly, although crRNA14 and crRNA15 target neighbouring regions, separated by just 8 nucleotides, their silencing efficiencies were markedly disparate. For example, 5ng of crRNA15 silenced >99% of mCherry expression (P<0.0001), while the same amount of crRNA14 did not show significant silencing of mCherry (P=0.78) (**Extended Data Figure 1c-1d**). As these two crRNAs target spatially adjacent sequences, this finding strongly suggested there are determinants of pspCas13b efficacy beyond target accessibility. Identifying such determinants is crucial in optimising crRNA design.

**Extended Data Figure 2.**
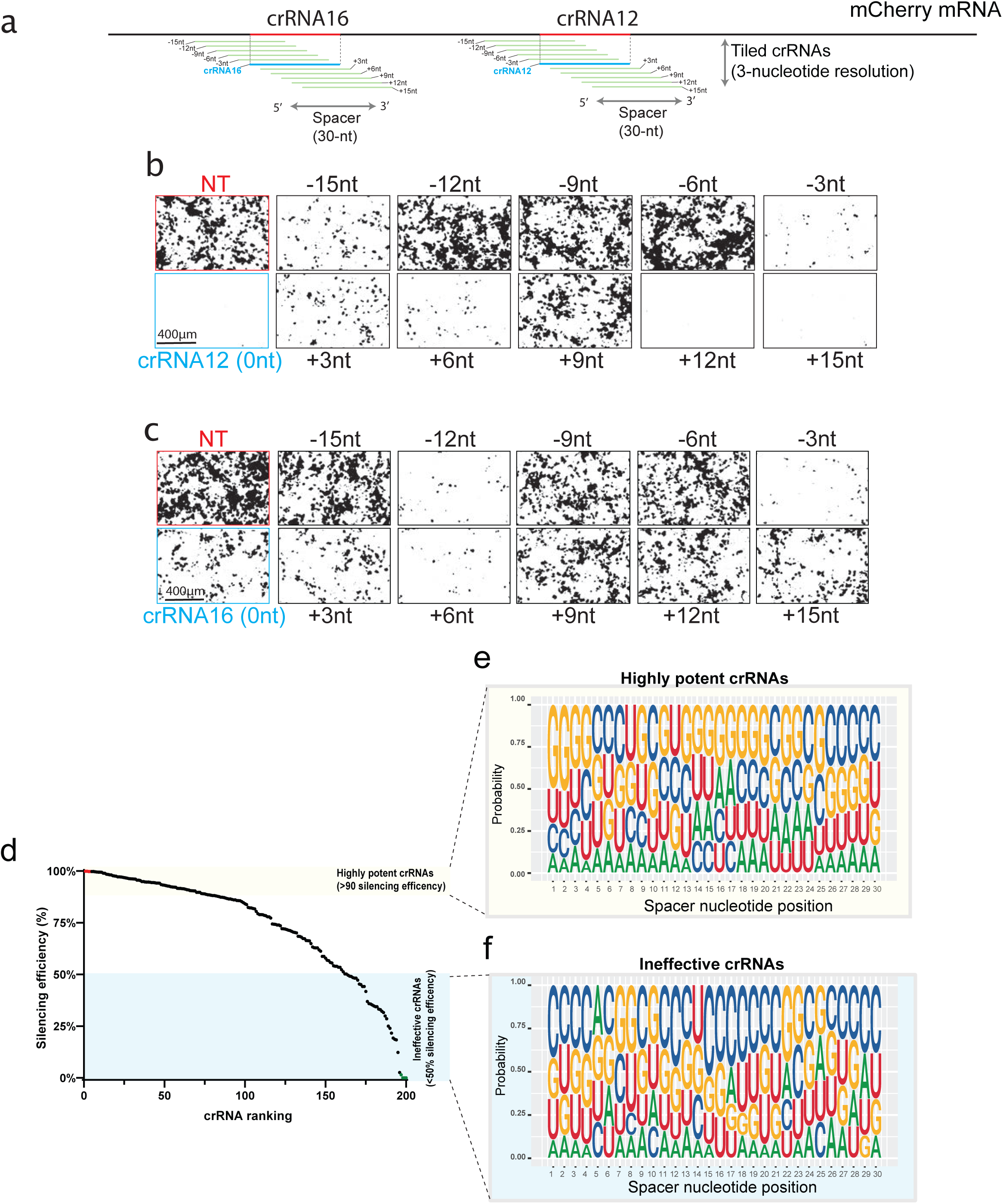
**(a)** The schematic illustrates mCherry RNA regions covered by 3-nucleotide increments tiled crRNAs. **(b-c)** Representative fluorescence microscopy images show the silencing of mCherry transcripts with tiled crRNAs targeting the region surrounding crRNA12 **(b)** and crRNA 16 **(c)**. Images are processed for quantification using ImageJ. Similar results were obtained in N=4. Unprocessed representative images are provided in the Source Data file. **(d)** crRNAs are ranked based on their silencing efficiency. The highly potent crRNAs that achieved >90% silencing efficiency (yellow) and the ineffective crRNAs that achieved <50% silencing efficiency (blue) are analysed for PFS and spacer nucleotide positions. **(e-f)** Position Weight Matrices (PWMs) depicting the positional nucleotide probabilities of either the highly potent **(e)** or the ineffective **(f)** crRNA spacer sequences.

**Extended Data Figure 3.**
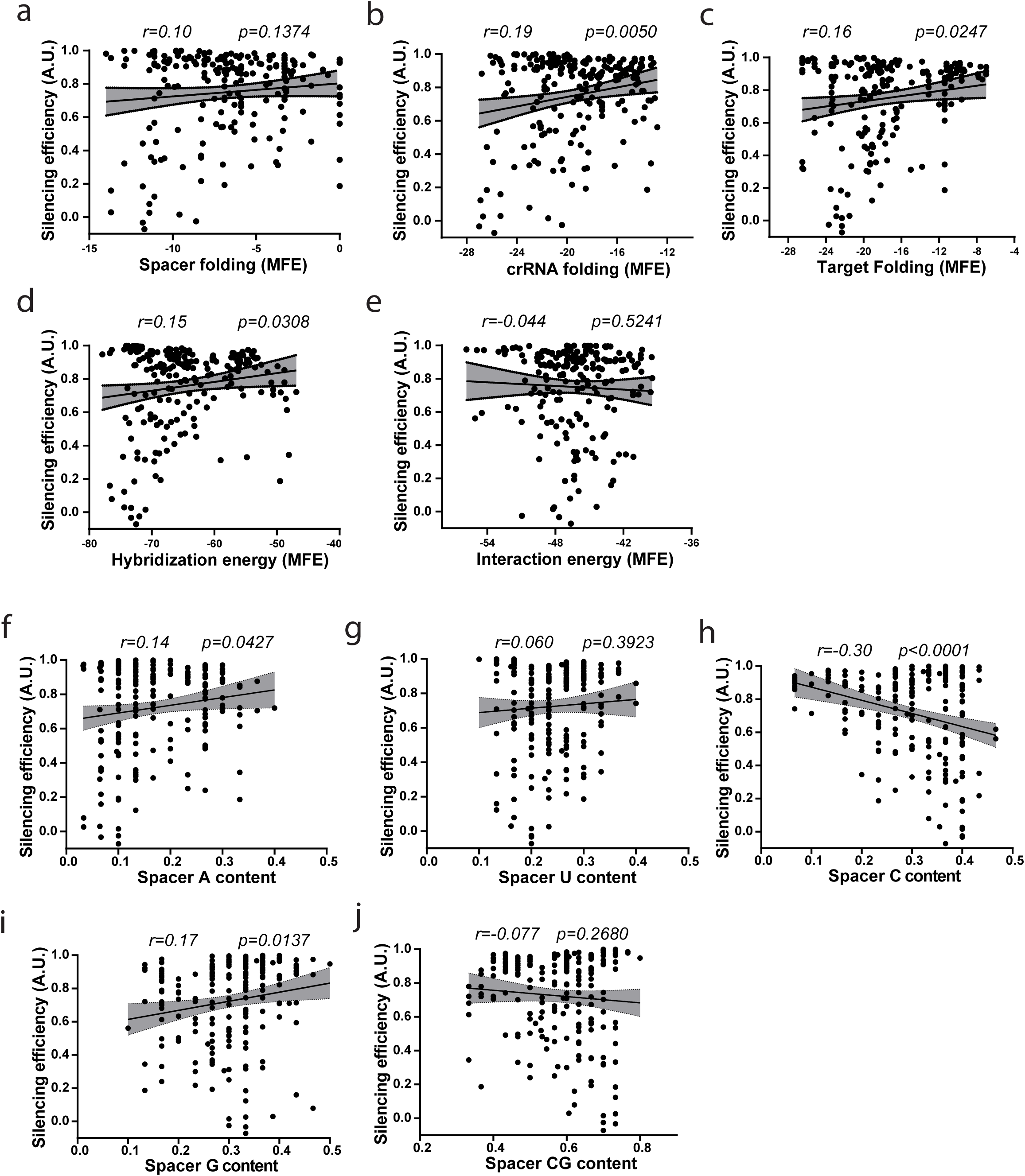
Pearson correlation analysis between **(a)** crRNAs silencing efficiency and spacer folding, **(b)** entire crRNA folding (spacer and direct repeat), **(c)** target sequence folding, **(d)** spacer-target hybridization, **(e)** and spacer-target interaction. Data points in the graph are values of the silencing efficiency of individual crRNAs and their predicted folding (MFE) or hybridization/interaction energy. Pearson correlation analysis between spacer silencing efficiency and the content of spacer in **(f)** A bases, **(g)** U bases, **(h)** C bases, **(i)** G bases, and **(j)** CG bases. Data points in the graph show the silencing efficiency and base content of individual spacer sequences. *r* (correlation coefficient) and *p-*value (95% confidence interval) are indicated in each graph. **Note:** We analysed a number of characteristics that may influence *Psp*Cas13b silencing efficiency in unfiltered crRNAs population including the predicted folding of crRNA, target folding, spacer-target hybridization energy (**Extended Data Figure 2a-2e**), and spacer nucleotide content (A, U, C, G, and CG) (**Extended Data Figure 2f-2j**). We used Pearson correlation to probe the existence of any relationship between the predicted folding and *Psp*Cas13b silencing efficiency. The data revealed a moderate positive correlation between the minimum free energy (MFE) of the crRNA and *Psp*Cas13b silencing efficiency (*r*=0.19; *p*=0.0050) (**Extended Data Figure 2b**). Similarly, target accessibility (folding of a 70-nt RNA sequence surrounding the targeted region) exhibited a modest correlation with the silencing efficiency of crRNA (*r*=0.16; *p*=0.0247) (**Extended Data Figure 2c**). Together, these data suggest that the folding of the crRNA and the targeted sequence into complex secondary structures can only moderately limit *Psp*Cas13b silencing efficiency, possibly perturbing crRNA loading or target accessibility. Next, we used a similar approach to analyse the effect of differential ribonucleotide abundance within the spacer on crRNA activity. A (*r*=0.14; *p*=0.0427) and G (*r*=0.17; *p*=0.0137) nucleotide content were positively correlated with *Psp*Cas13b silencing, whereas spacers with enriched C content exhibited a strong negative correlation with crRNA potency (*r*=-0.30; *p*<0.0001) (**Extended Data Figure 2f-3j**). These data indicate that spacer nucleotide content is likely a key determinant of *Psp*Cas13b silencing, and suggest spacer enrichment with C bases is likely to impede their potency.

**Extended Data Figure 4.**
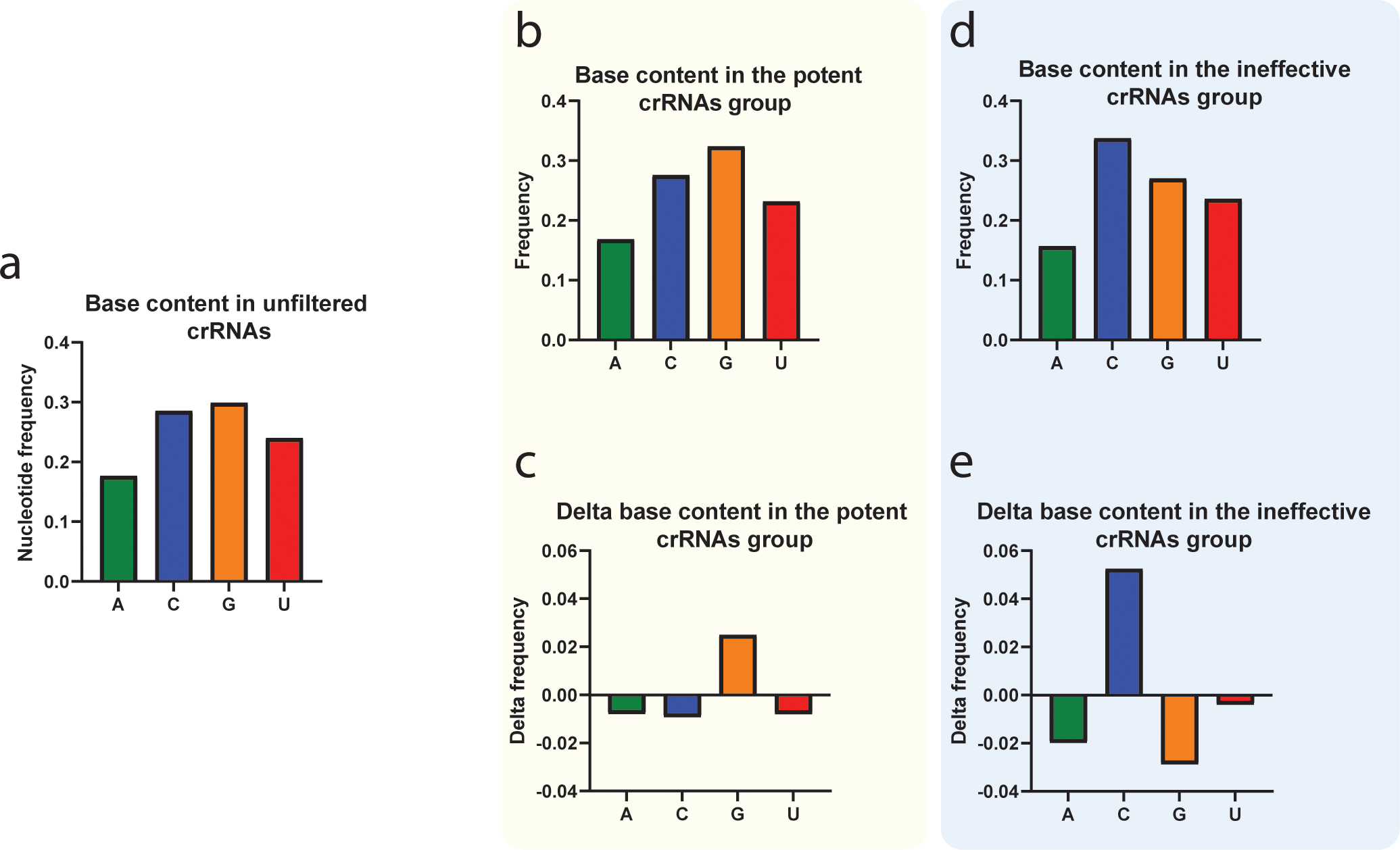
**(a)** The frequency of A, C, G, and U bases in unfiltered crRNA spacer sequences. The frequency **(b)** and delta frequency **(c)** of each base in the spacer sequences of filtered potent crRNAs. The frequency **(d)** and delta frequency **(e)** of each base in the spacer sequences of filtered ineffective crRNAs. **Note:** Overall, G nucleotides are enriched in the spacer sequences of potent crRNAs compared to the unfiltered total crRNAs (**Extended Data Figure 4a-4c**). Conversely, C nucleotides are enriched in the ineffective crRNA cohort (**Extended Data Figure 4a, 4d, & 4e**). This data reveals that potent crRNA favours G over C bases, although the positions of the enriched/depleted G or C nucleotides in the spacer of effective and ineffective crRNAs remain unknown.

**Extended Data Figure 5.**
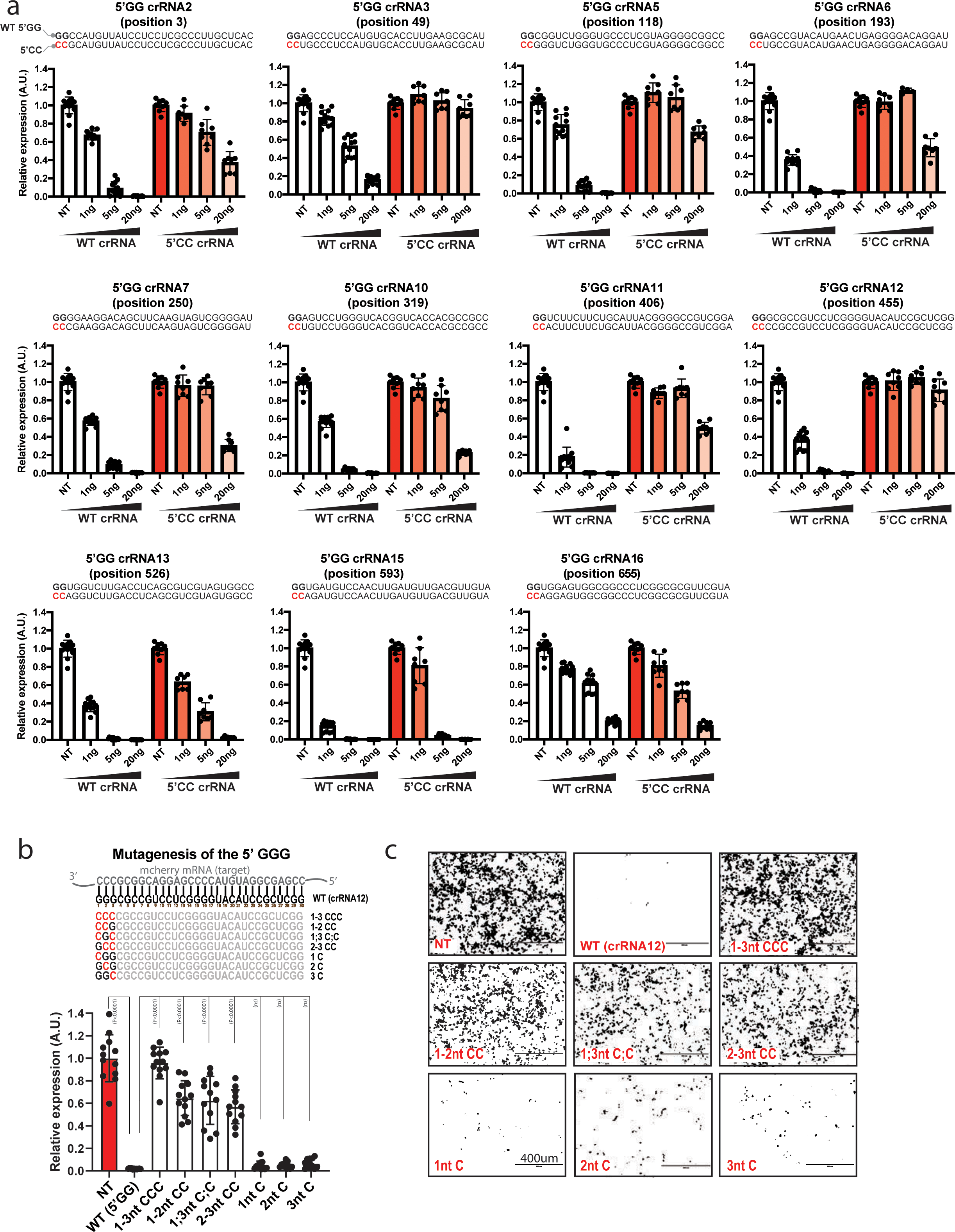
**(a)** Dose-dependent silencing of mCherry transcript with non-targeting crRNA (NT), 11 unmodified crRNA (WT) that possess a GG sequence at their 5’end (white bars), or the same 11 crRNAs were the 5’ GG sequence of the spacer is mutated to 5’ CC through 1-3 nucleotides mutagenesis. Data points in the graph are normalized mean fluorescence from 4 different field of views imaged in *N*=2. The data are represented in arbitrary units (A.U.). Errors are SD with 95% confidence interval. **(b)** Mutagenesis analysis of spacer 1-3 nucleotides (5’ end) examining the impact of C to G substitutions on crRNA silencing efficiency. The nucleotides in red highlight mismatch positions in the spacer sequence. Data points in the graph are mean fluorescence from 4 representative field of views per condition imaged; Errors are SD and p-values of one-way Anova test are indicated (95% confidence interval). *N* = 3. The data are represented in arbitrary units (A.U.). *N* is the number of independent biological replicates. Source data are provided as a Source data file. **(c)** Representative fluorescence microscopy images show the silencing efficiency of the mCherry transcripts with NT, WT and mutant crRNAs in HEK 293T cells. NT is a non-targeting control crRNA. Scale bar = 400μm. Similar results were obtained in 3 independent experiments in HEK 293T cells. Unprocessed representative images are provided in the Source Data file. **Note:** The dose-dependent silencing assays show that the substitution of G to C bases at the 5’ end of the spacer greatly compromised their silencing efficiency. For instance, 5 ng of WT crRNA7 achieved ∼95% silencing, whereas the substitution of GG by a CC sequence led to a complete loss of silencing (**Extended Data Figure 5a**). The data also show that two out of three first nucleotides at the 5’end of the spacer are required to be G bases to preserve crRNA potency for this spacer (**Extended Data Figure 5b & 5c**).

**Extended Data Figure 6.**
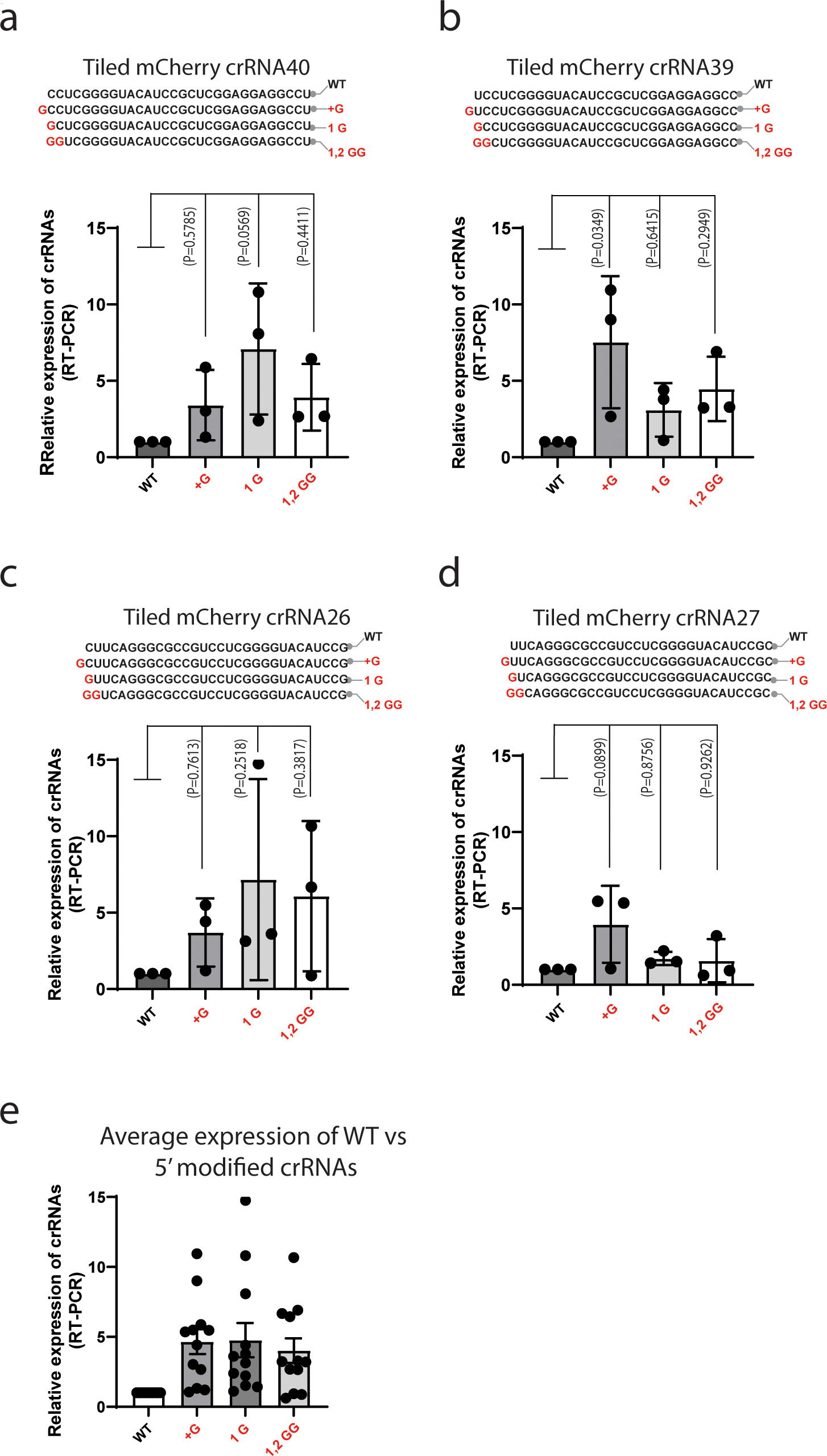
**(a-d)** RT-PCR analysis to assess the expression of WT crRNAs, crRNAs with an extra G at their 5’ end (31-mer spacer), crRNAs with the first spacer nucleotide substituted to a G (30-mer spacer), and crRNAs with the first and second spacer nucleotides substituted to GG (30-mer spacer). crRNAs expression was measured 48h post-transfection in HEK 293T cells, *N* = 3. Data are normalized means and errors are SEM; Results analysed with one-way Anova test with *p-*value indicated (95% confidence interval). **(e)** Averaged expression (from a-d) of unmodified or modified crRNAs harbouring G-rich 5’end. **Note:** The data indicate that the incorporation of a G-rich motif at the 5’end (first and second positions) of the spacer increases crRNA expression or stability. Previous study ^1^ suggested that U6 promotor has a preference for a G or A nucleotides surrounding the transcription starting site. Therefore, systematic design of crRNA with two G nucleotides at the 5’ end may increase crRNA potency through enhanced transcription.

**Extended Data Figure 7.**
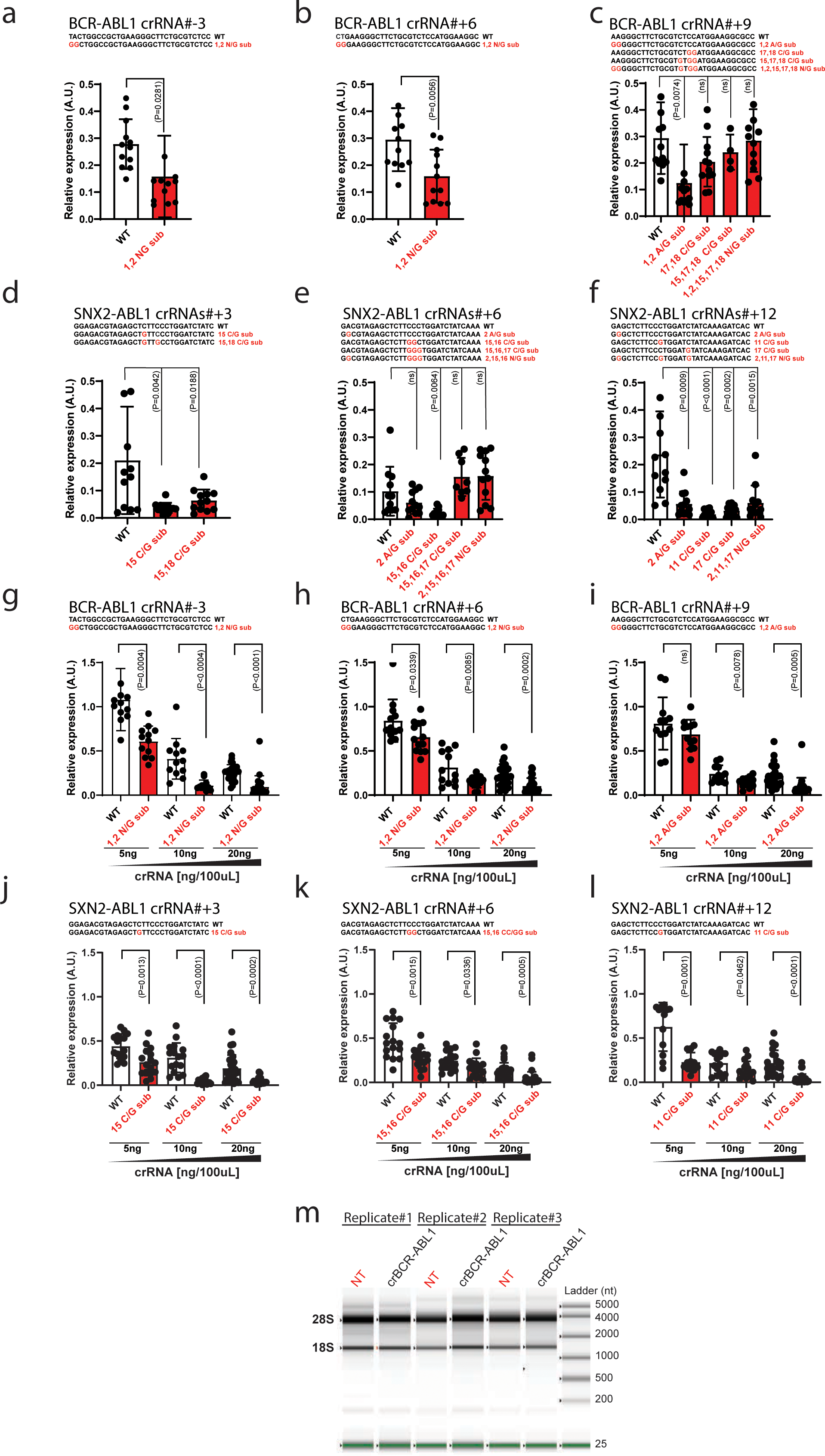
Incorporation of target-mismatched ‘G’ bases at the 5’end of spacer sequence greatly enhances *Psp*Cas13b crRNA efficiency. (**a-f**) Comparison of silencing efficiencies of crRNAs targeting the breakpoint of gene fusion transcripts with or without incorporation of mismatched G-bases at the 5’end and/or central regions of the spacer. Data points in the graphs are mean fluorescence from 4 representative field of views per condition imaged; *N* =3 or 4. The data are represented in arbitrary units (A.U.). Errors are SD and *p*-values of unpaired two-tailed Student’s t-test are indicated (95% confidence interval). **(g-l)** crRNA dose-dependent silencing of gene fusion transcripts with or without incorporation of mismatched G-bases at the 5’end and/or central regions of the spacer. Data points in the graphs are mean fluorescence from 4 representative field of views per condition imaged; *N* =3 or 4. The data are represented in arbitrary units (A.U.). Errors are SD and *p*-values of unpaired two-tailed Student’s t-test are indicated (95% confidence interval). Data points in the graphs are mean fluorescence from 4 representative field of views per condition imaged; *N* =3 or 4. *N* is the number of independent biological replicates. Source data are provided as a Source data file. (**m**) Assessment of potential *Psp*Cas13b collateral activity through the analysis of ribosomal RNA (28S and 18S) degradation in the presence of target-free (NT) or target-bound (crBCR-ABL1) *Psp*Cas13b nuclease. **Note**: The silencing assays in **Extended Data Figure 7a-7l** show that the substitution of nucleotides at spacer positions 1 and 2 with a GG sequence enhances the potency of six different crRNAs targeting gene-fusion transcripts. Likewise, the substitution of ‘C’ base at the central region (positions 11, 12, 15, 16, 17, 18) with a G base also enhance the potency of six crRNAs targeting the breakpoint of oncogenic gene fusion transcripts BCR-ABL-1 and SNX2-ABL1. These data confirm the importance of 5’ G-rich motif and the avoidance of C bases at the central region (positions 11, 12, 15, 16, 17, 18) for maximum silencing efficiency, reinforcing the data described in Figure 1 **&** Figure 2. The incorporation of up to 3 mismatched G bases per spacer (5’end and/or central region) can enhance the potency of crRNAs, but beyond 3-nucleotide substitutions, the benefit of mismatched G bases is often compromised (**Extended Data Figure 7c, 7e**) likely due to the destabilization of spacer-target interaction, in agreement with mutagenesis data in **Extended Data Figure 8** and Figure 4a**, 4c**. The data in **Extended Data Figure 7m** suggests that the silencing of BCR-ABL1 fusion transcript with *Psp*Cas13b does not lead to a global degradation of RNA due to its collateral activity, as evidenced by the stability of ribosomal RNAs 28S and 18s regardless of the use of NT or BCR-ABL1 targeting crRNAs. This data suggests high on-target activity of *Psp*Cas13b with no or very minor collateral activity of *Psp*Cas13b. In contrast to these *Psp*Cas13b data, target recognition and cleavage of *Rfx*Cas13b has been shown to trigger collateral degradation of various host cellular RNAs including the 28S ribosomal RNAs^2, 3^.

**Supplementary file 1.**
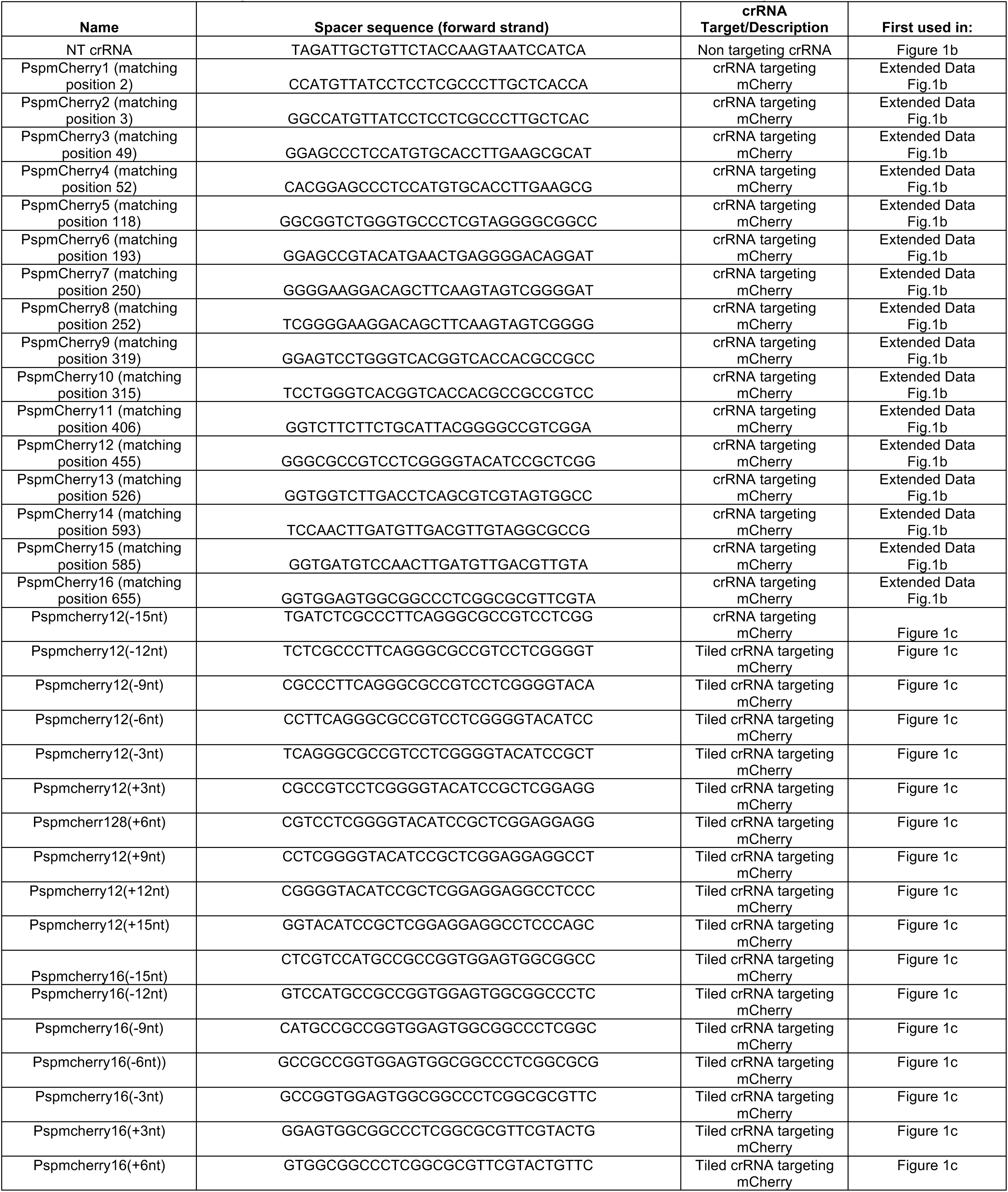
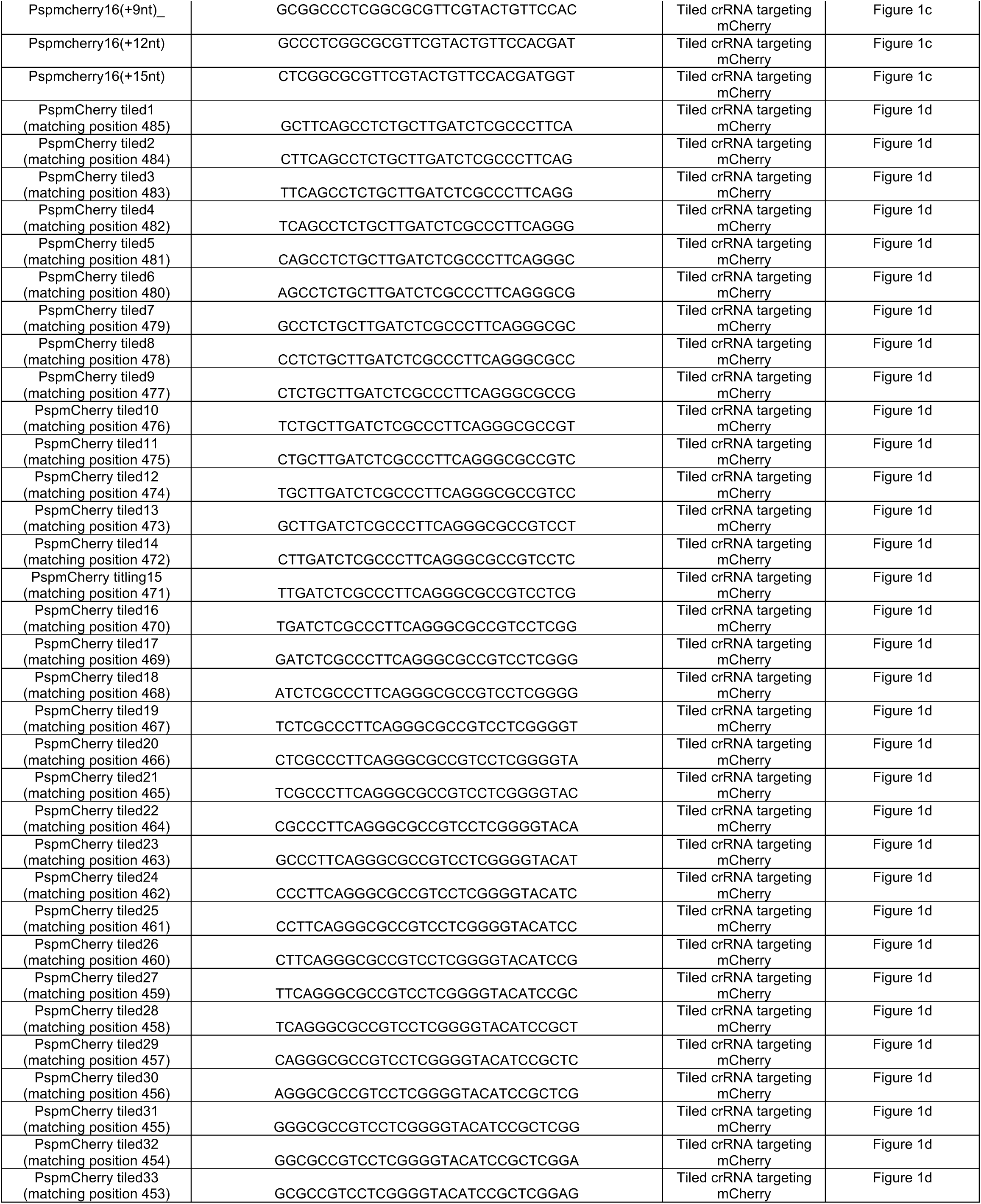
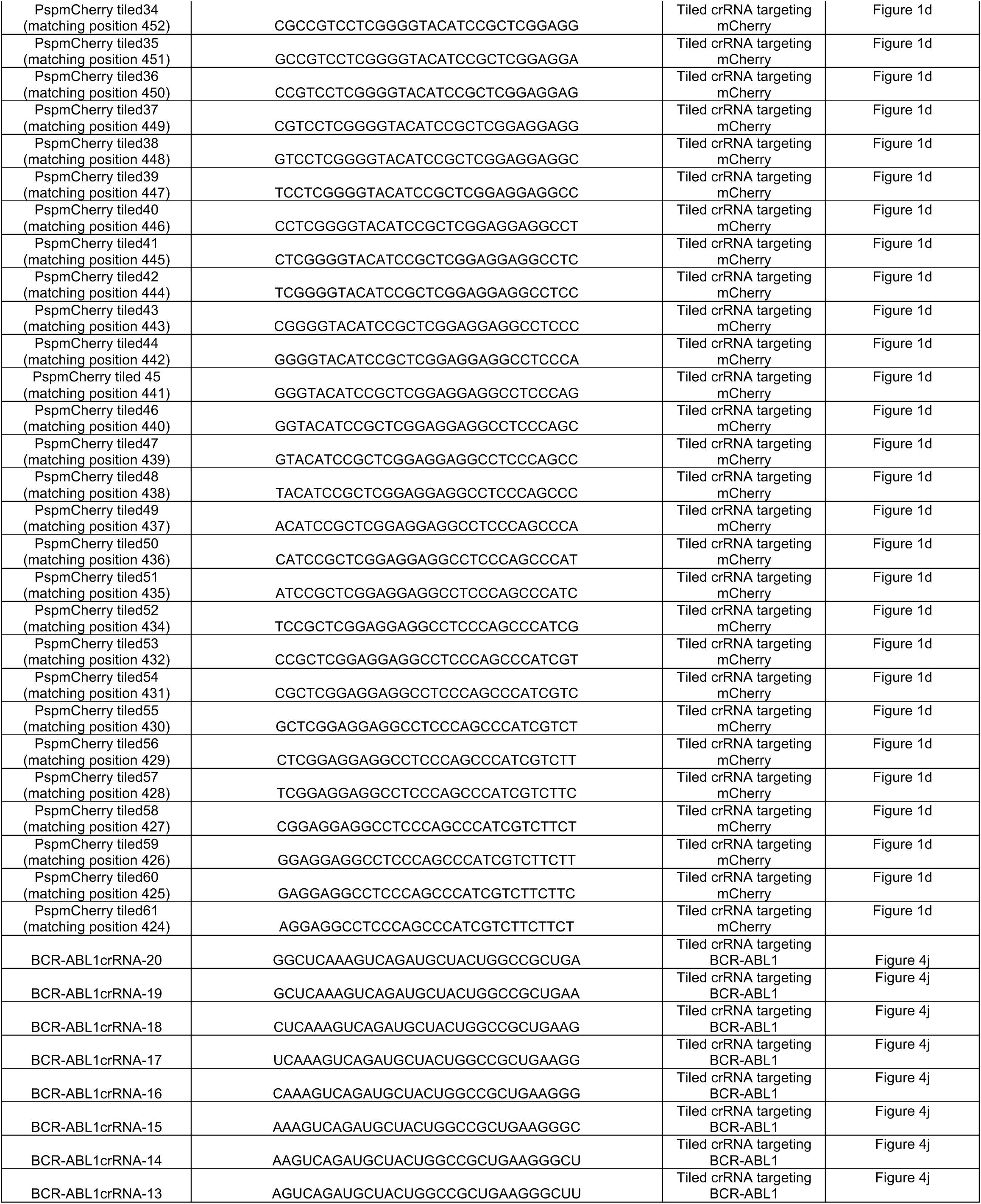
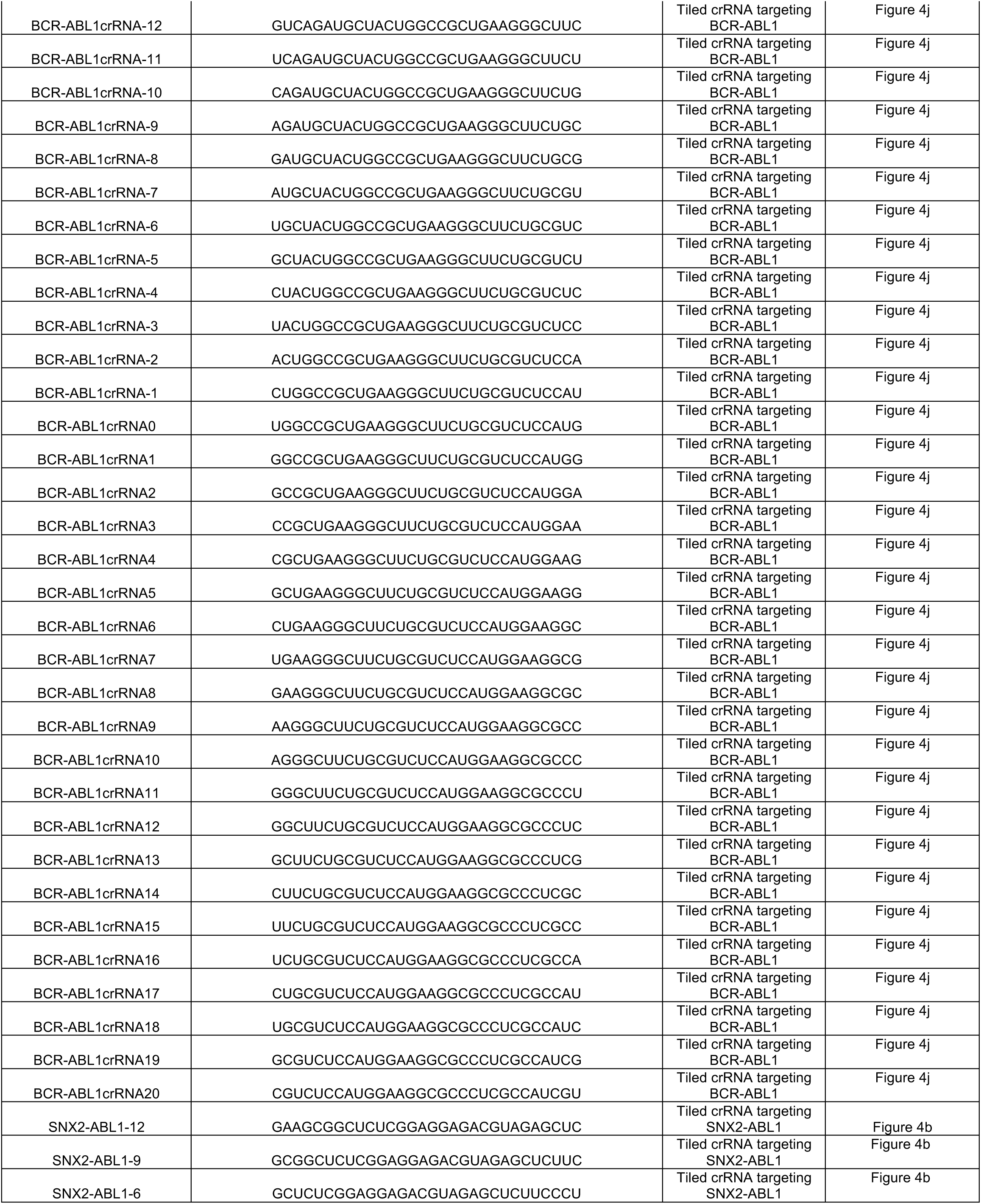
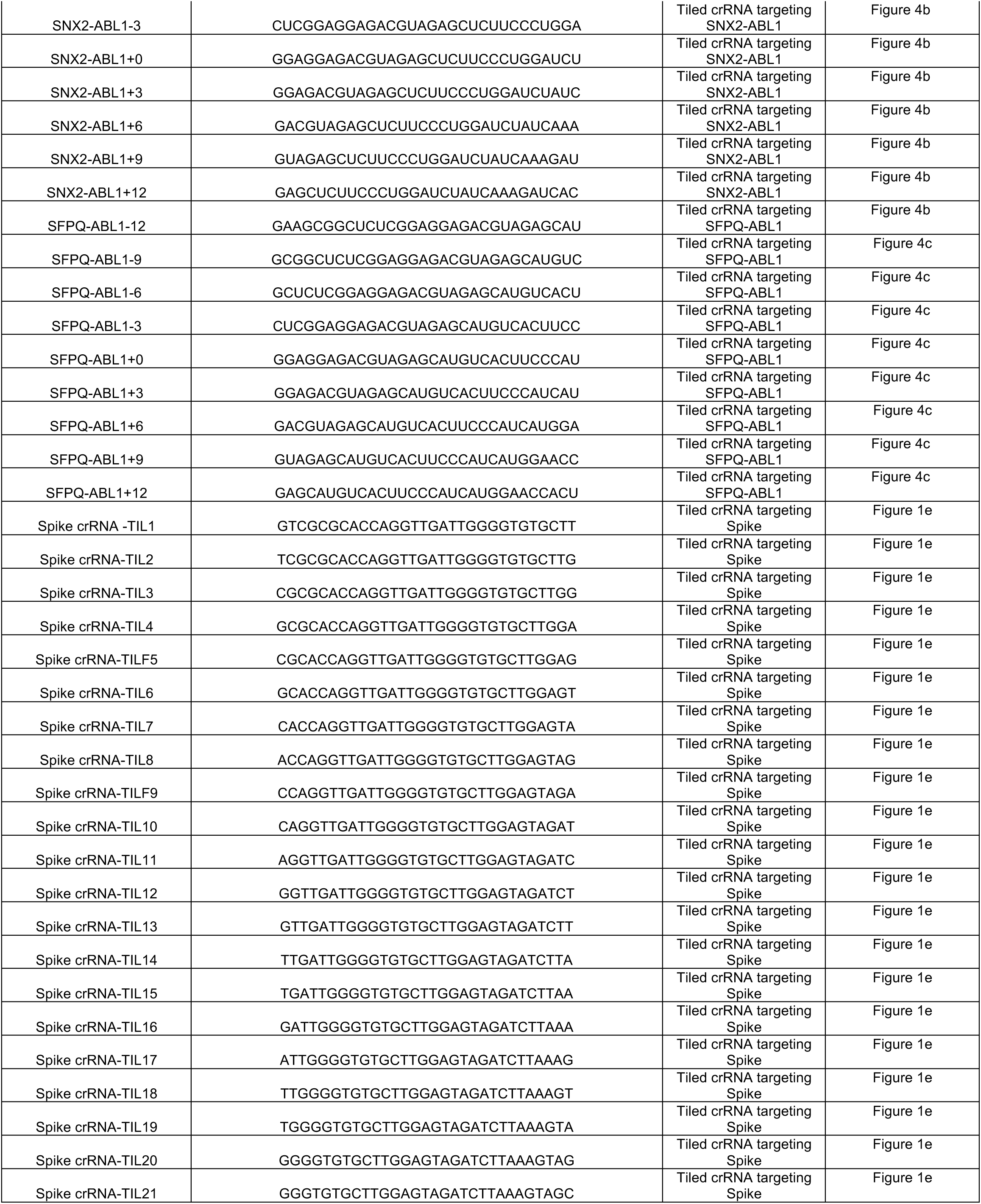
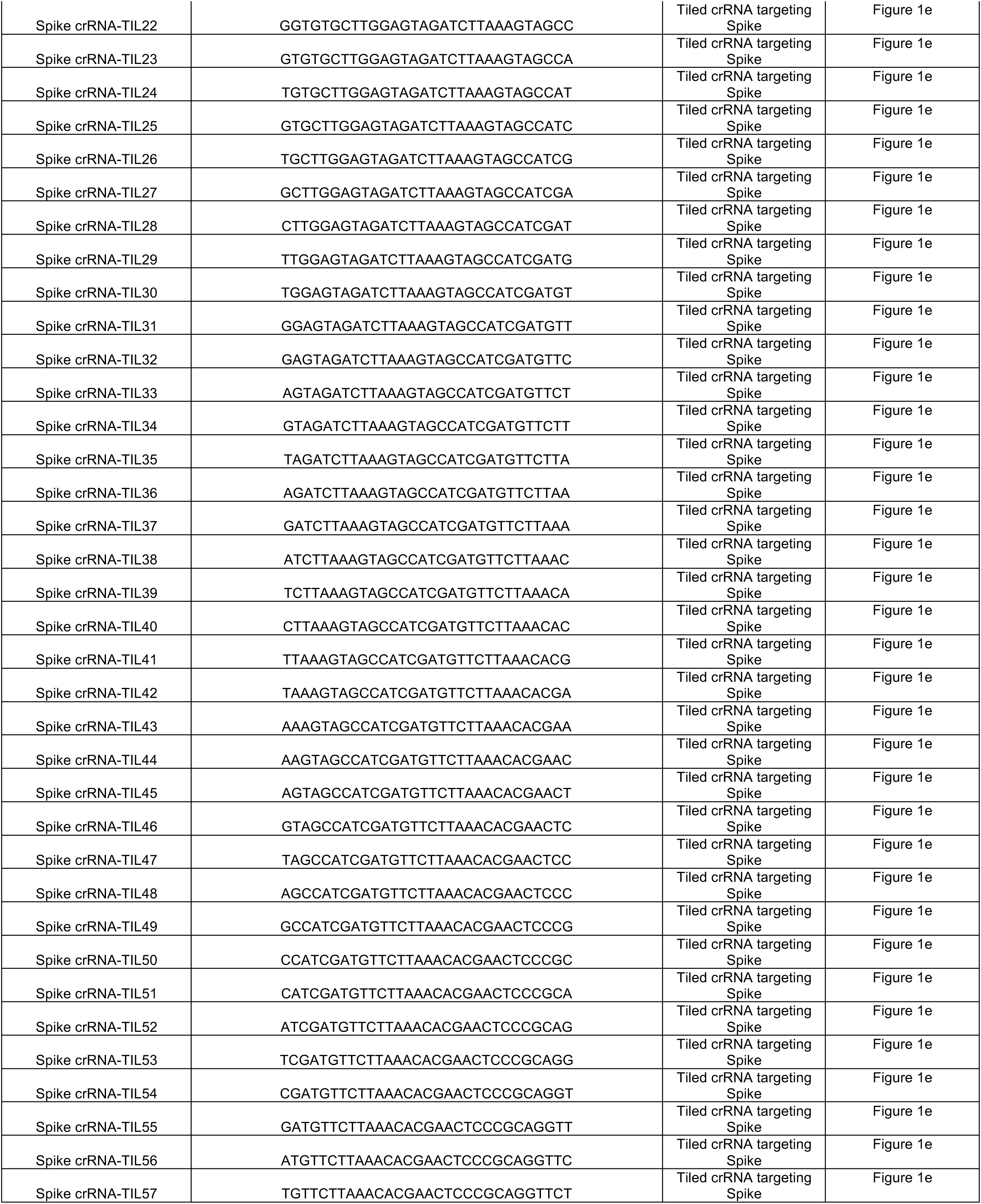
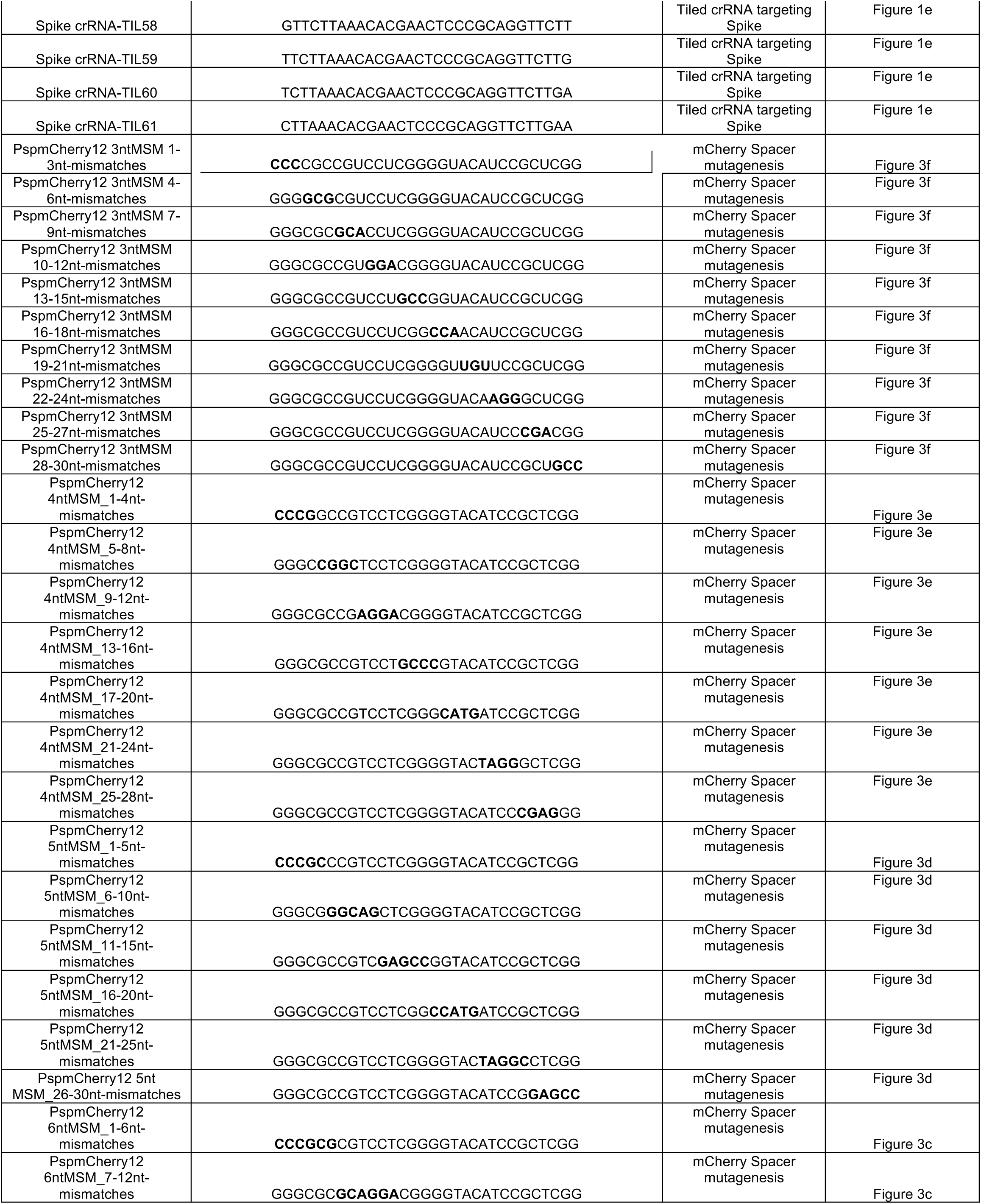
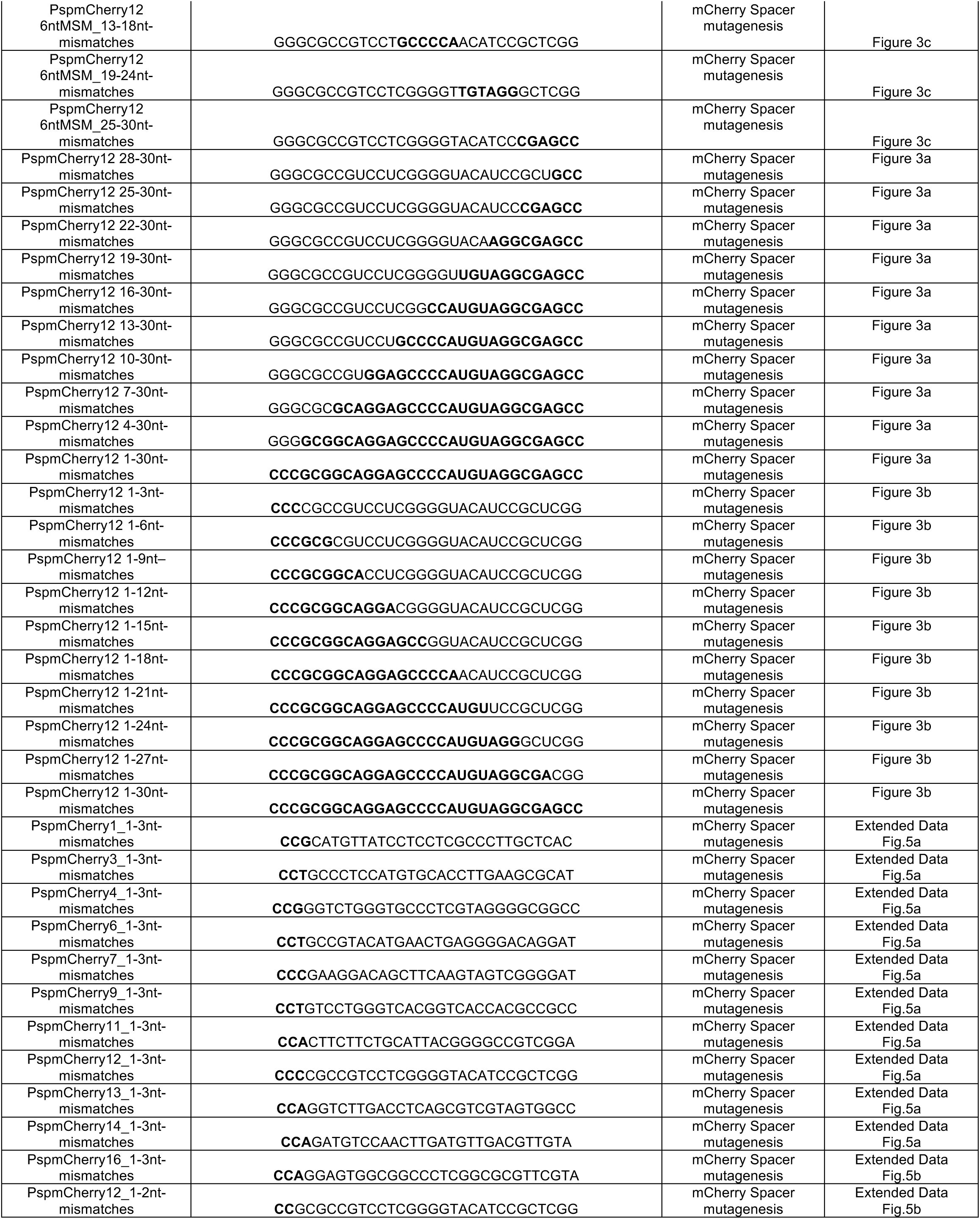
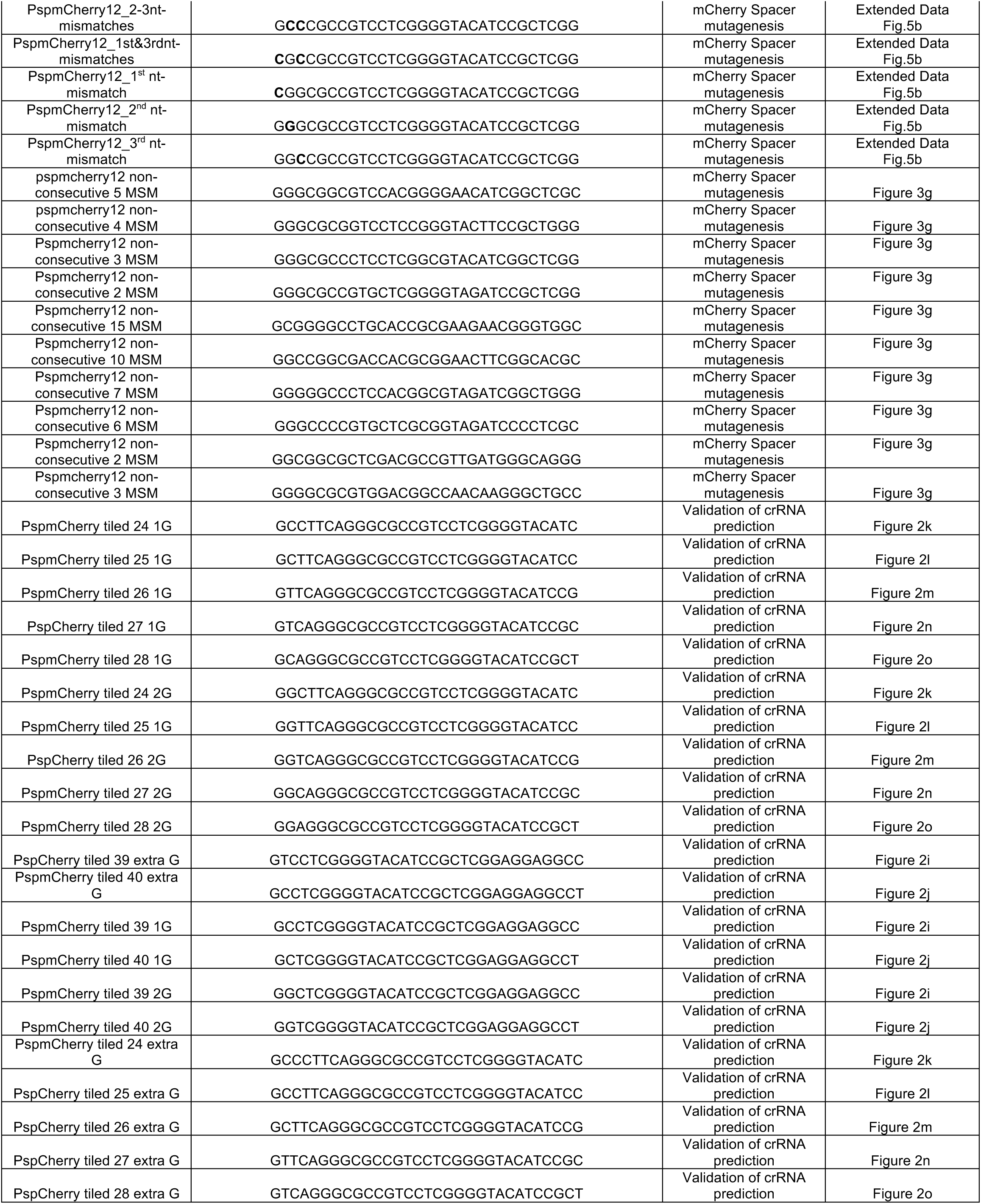
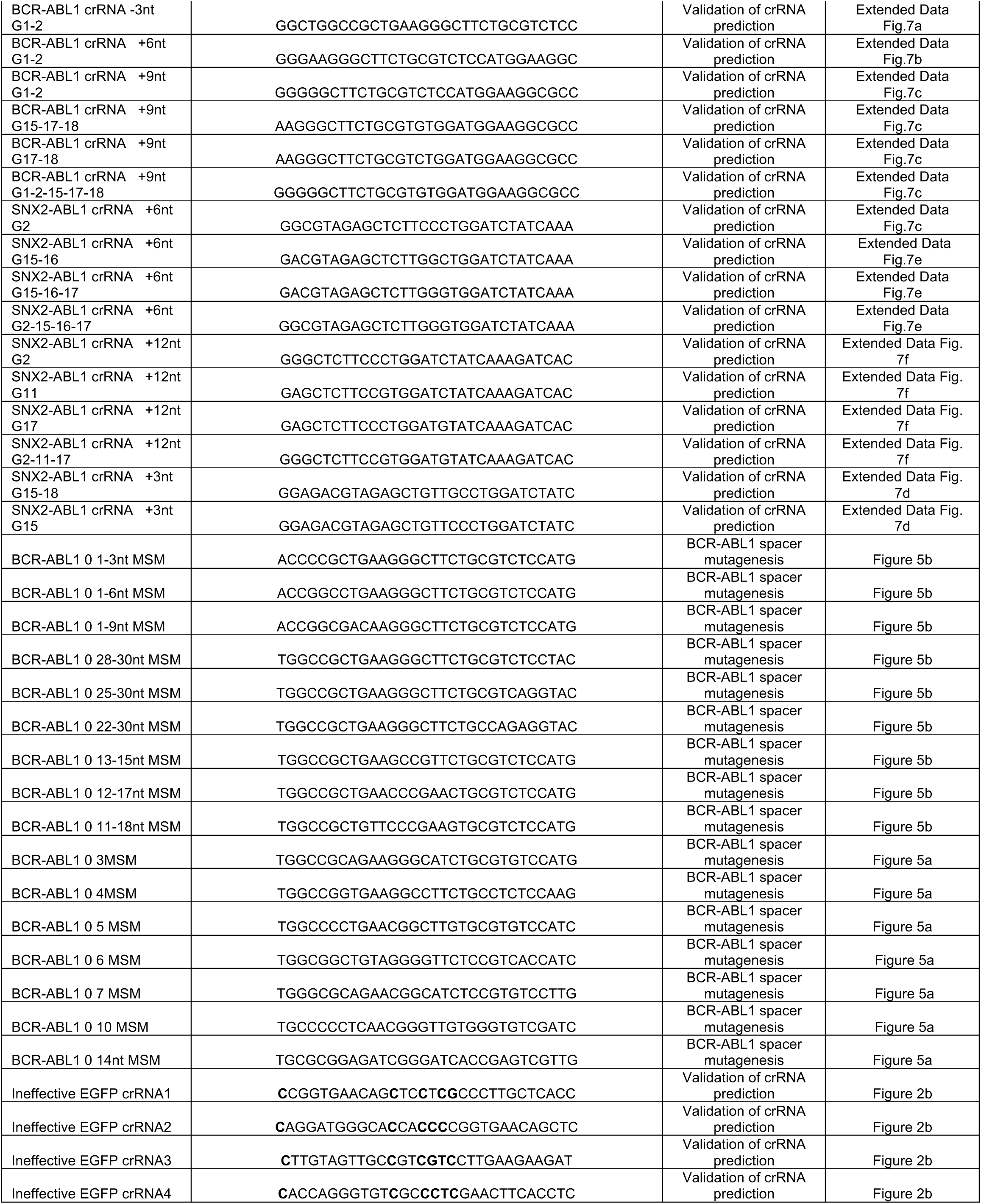
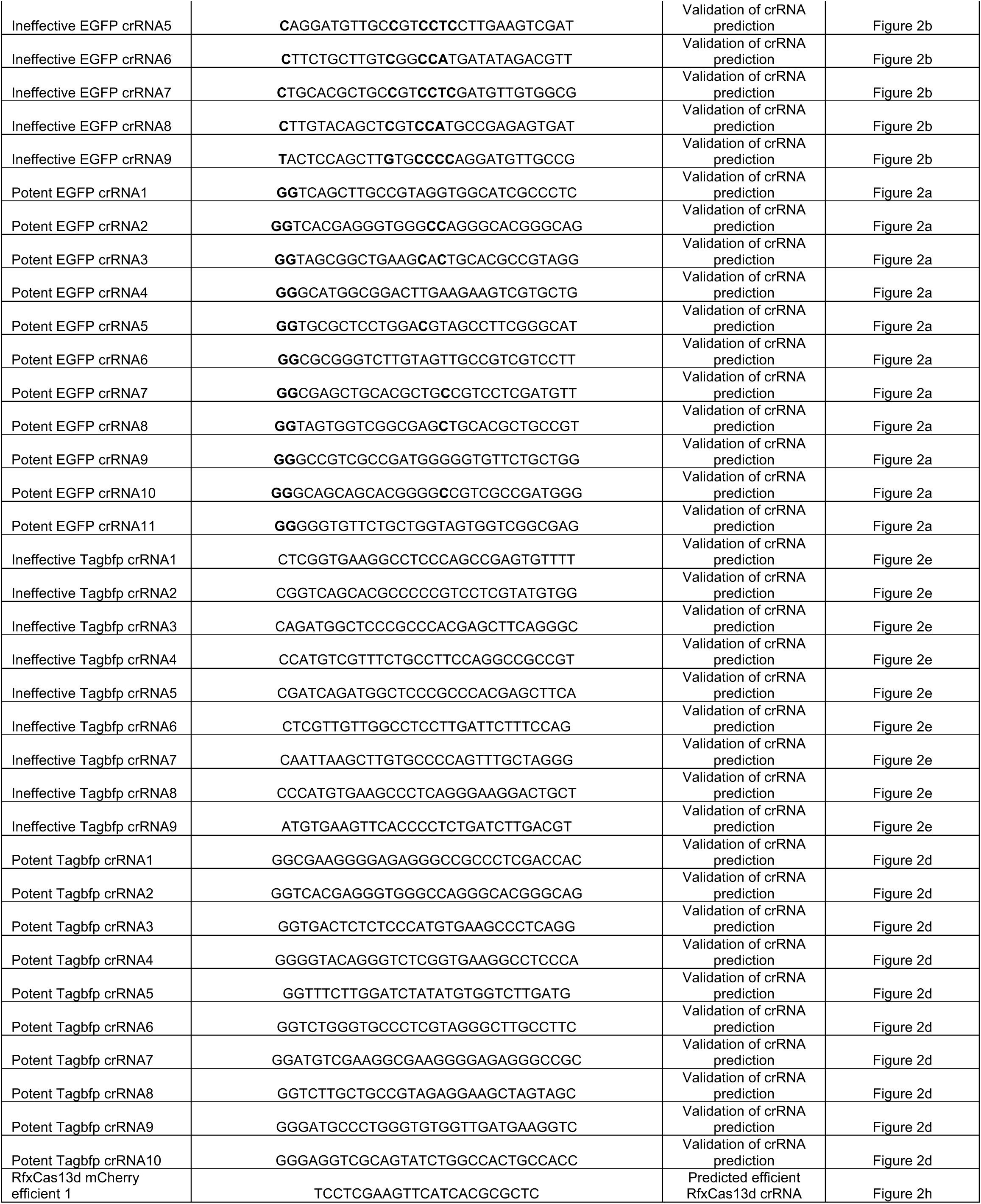
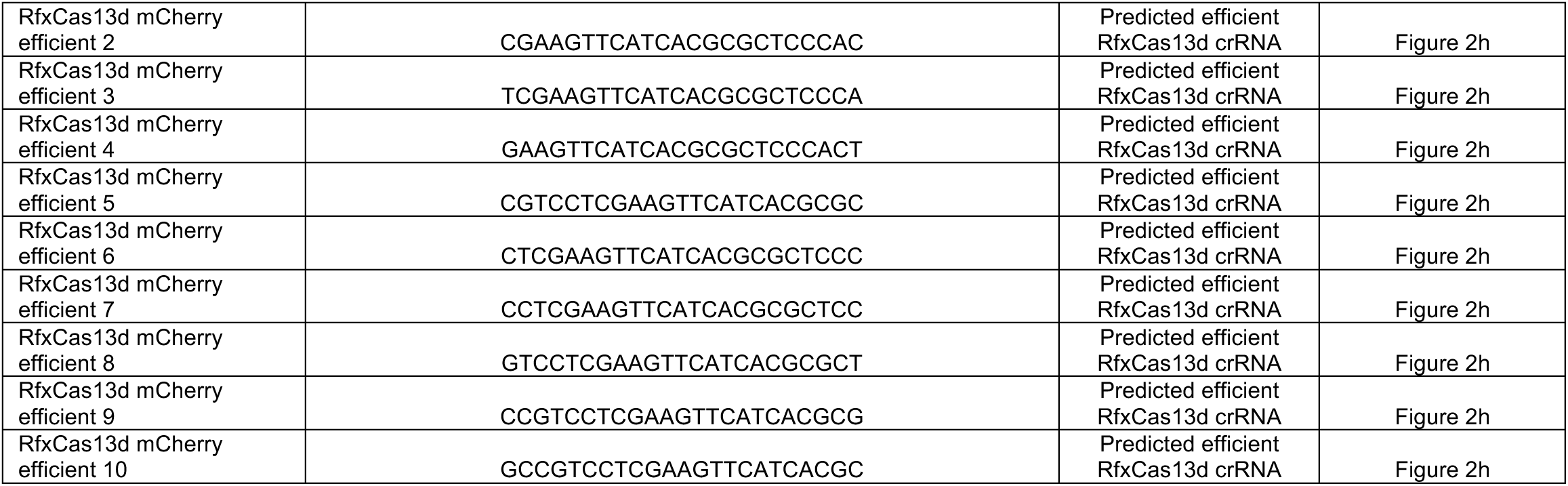
crRNA spacer sequences used in this study.

**Supplementary file 2.**
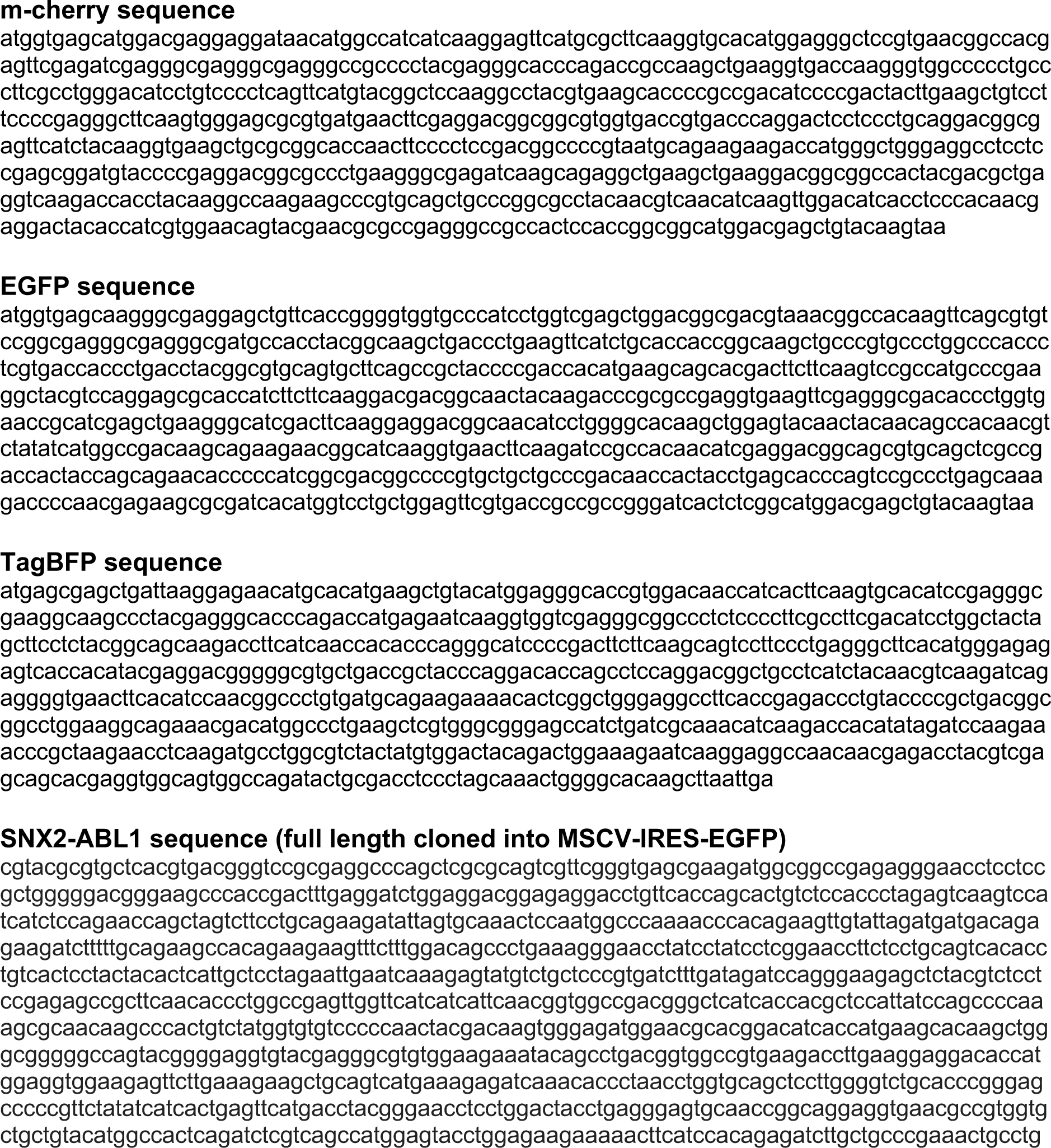
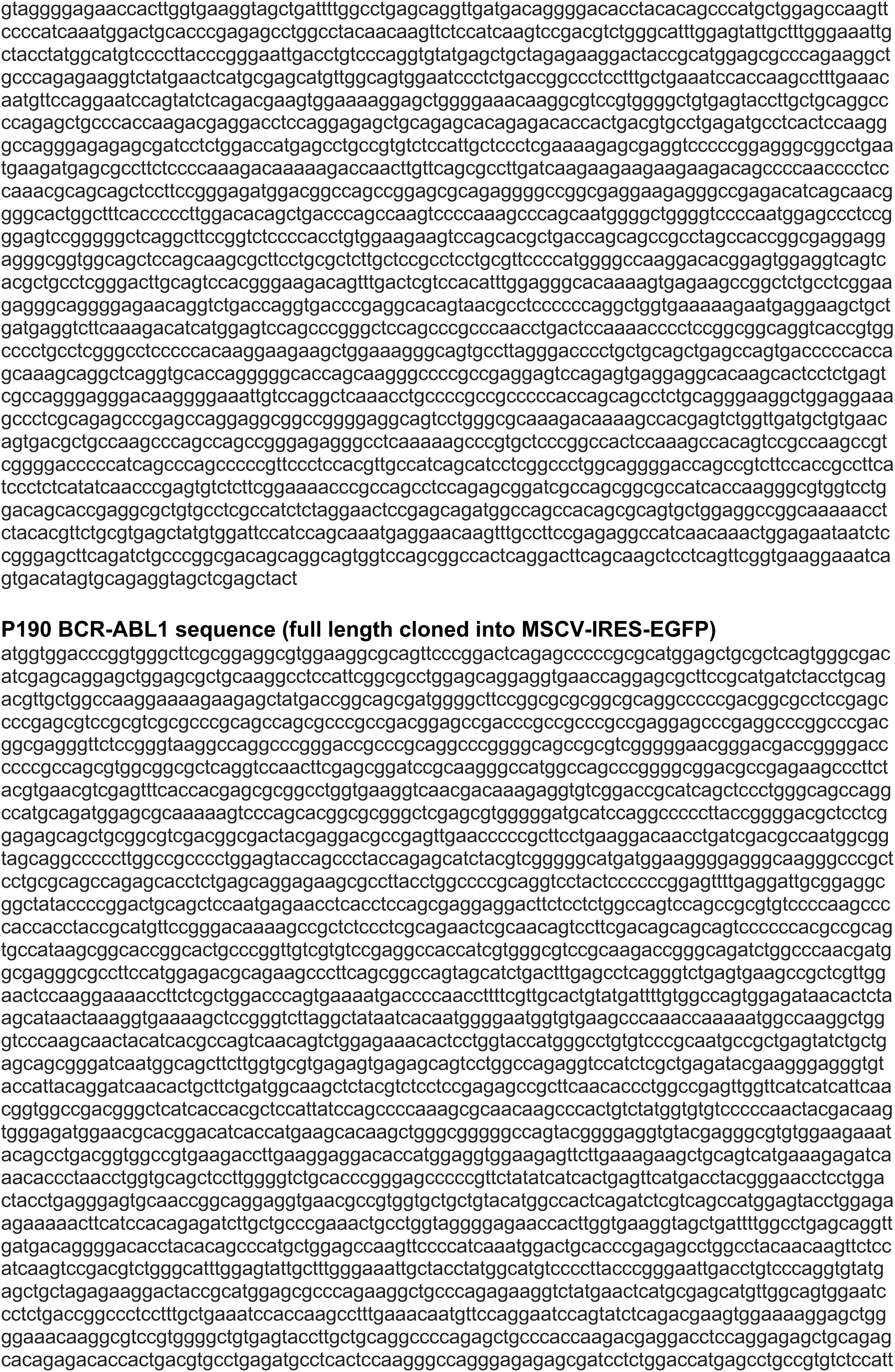
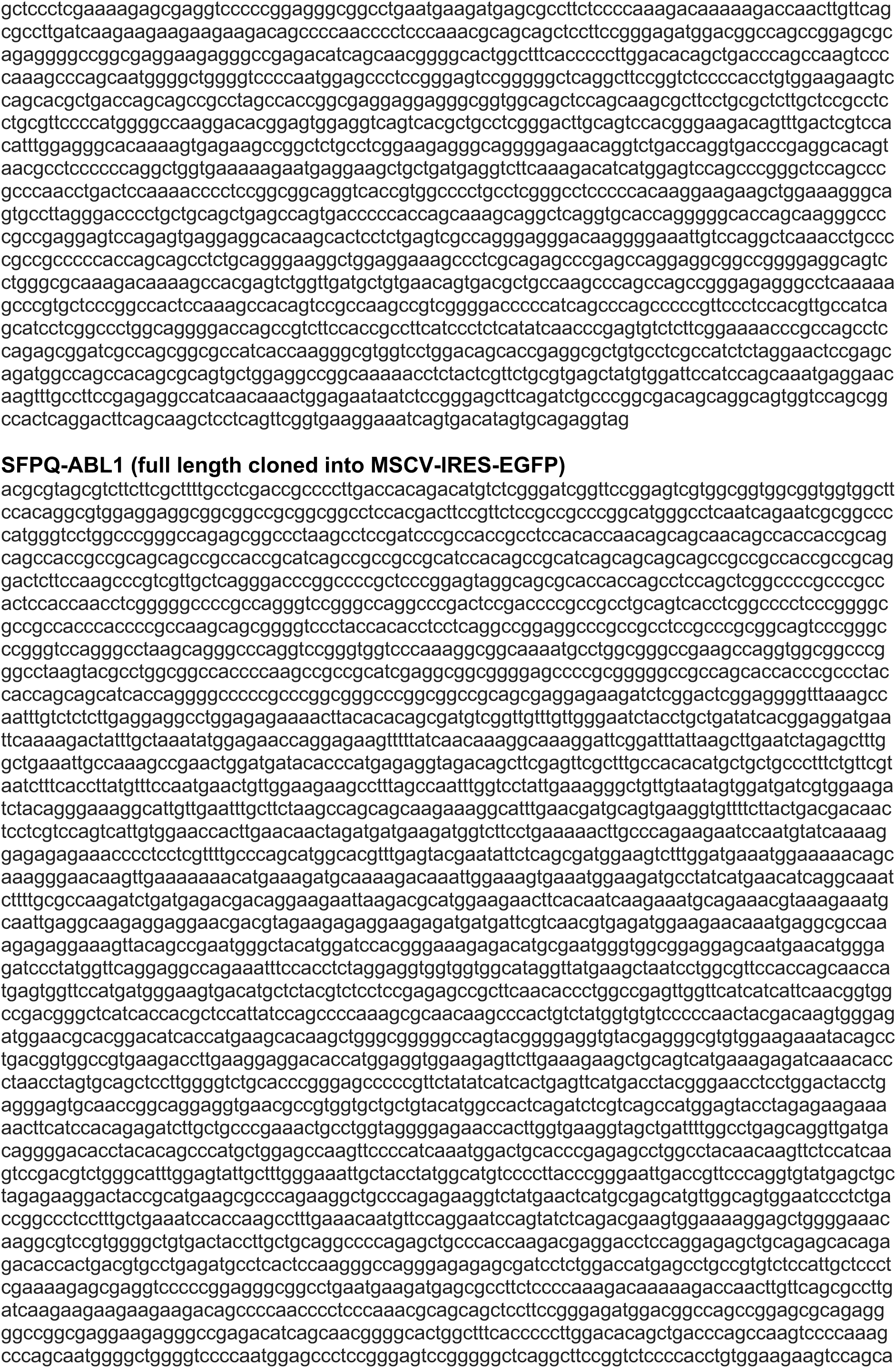
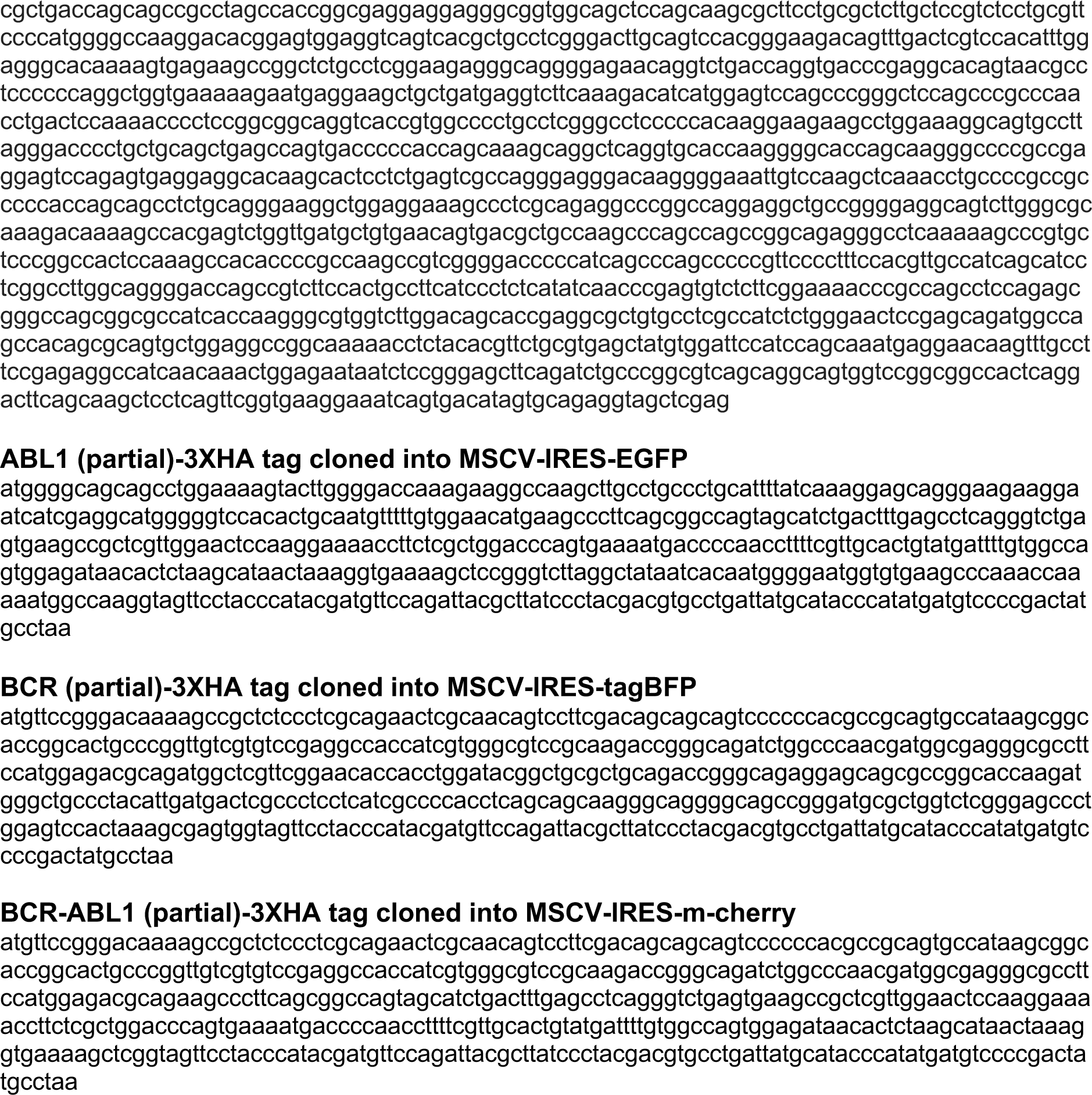
Target sequences used in this study.

**Supplementary file 3.**
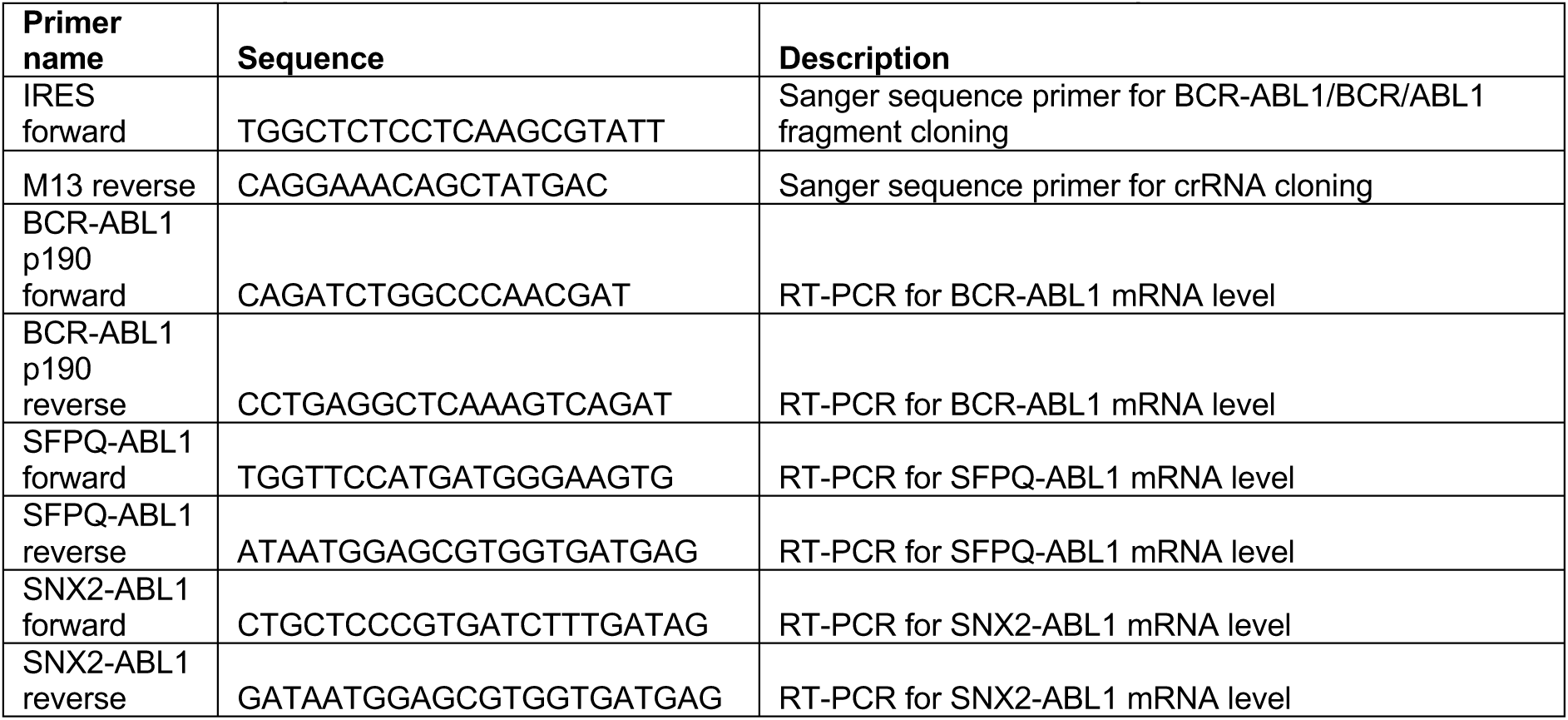
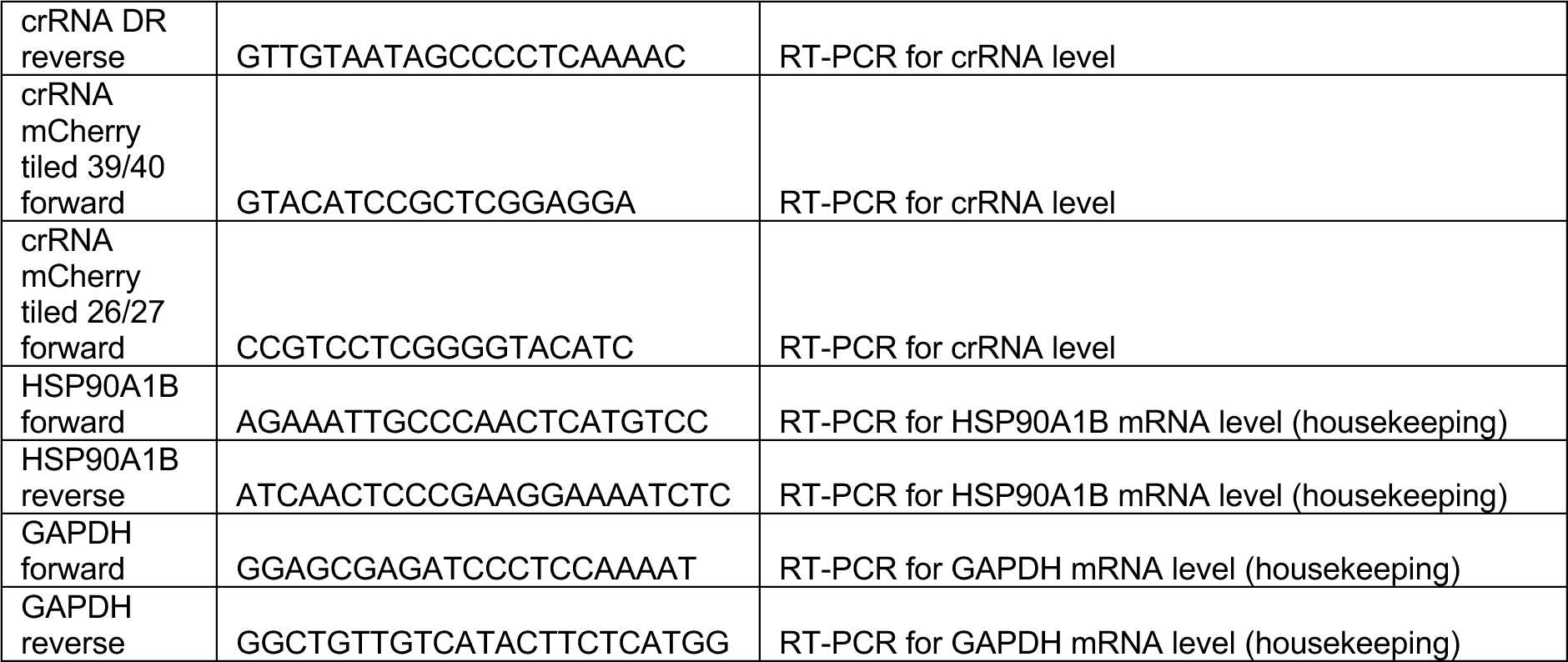
Primer sequences used in this study.

**Supplementary file 4.** Transfection conditions of HEK 293T cells.

| Component (per well) | 96-well | 24-well | 12-well |
| --- | --- | --- | --- |
| # of seeded HEK 293T cells | 30,000 | 150,000 | 300,000 |
| Plasmids DNA amount | 100ng | 500ng | 1000 ng |
| P3000 reagent | 0.2µL | 1µL | 2 µl |
| Lipofectamine 3000 reagent | 0.3µL | 1.5µL | 3 µl |
| Opti-MEM | Up to 10 µL | Up to 50 µL | Up to 100 µL |

**Supplementary file 5.** Resources Table.

| Reagent or resource | Source | Identifier |
| --- | --- | --- |
| **General chemicals** |  |  |
| Sodium deoxycholate | Sigma-Aldrich | D6750 |
| Ethanol | Sigma-Aldrich | 443611 |
| Methanol | Sigma-Aldrich | 34860 |
| Glycine | Sigma-Aldrich | G7126 |
| Isopropanol | Sigma-Aldrich | 563935 |
| Agarose | Sigma-Aldrich | A9539 |
| Sodium dodecyl sulphate (SDS) | Sigma-Aldrich | L3771 |
| Bovine serum albumin (BSA) | Sigma-Aldrich | A3059 |
| NP-40 | Sigma-Aldrich | I8896 |
| Tris | Sigma-Aldrich | T1530 |
| Tween-20 | Sigma-Aldrich | P1379 |
| Triton X-100 | Sigma-Aldrich | T8787 |
| **Western blot reagents** |  |  |
| Precision Plus Protein Dual Color Standards | Bio-Rad | 1610374 |
| PageRuler Plus Prestained Protein Ladder | Thermo Fisher | 26619 |
| Pierce BCA Protein Assay Kit | Thermo Fisher | 23225 |
| X4 Blot LDS sample buffer | Thermo Fisher | B0007 |
| X10 Bolt sample reducing agent | Thermo Fisher | B0009 |
| Bolt Bis-Tris Plus 4-12% gel | Thermo Fisher | NW04120BOX |
| 4-15% Criterion TGX Precast Midi Protein gels | Bio-Rad | 5671084 |
| 20x MES SDS running buffer | Thermo Fisher | B0002 |
| 10x Tris/glycine/SDS running buffer | Bio-Rad | 1610732 |
| ECL Select Western Blotting Detection Reagent | Cytiva Life Sciences | 12393969 |
| Complete mini protease inhibitor cocktail tablet | Roche | 4906845001 |
| protease inhibitor cocktail | Roche | 04693159001 |
| 0.45 μM Polyvinylidene fluoride (PVDF) membranes | Thermo Fisher | 88518 |
| **Constructs** |  |  |
| pC0046-EF1a-PspCas13b-NES-HIV | Addgene | 103862 |
| PspCas13b-NES-HIV-T2A-BFP | Addgene | 173029 |
| pC0043-PspCas13b crRNA backbone | Addgene | 103854 |
| pMSCV-IRES-m-Cherry | Addgene | 52114 |
| MSCV-IRES-EGFP | A gift from Ilia Voskoboinik Lab | N/A |
| MSCV-IRES-tagBFP | A gift from Ilia Voskoboinik Lab | N/A |
| MSCV-BCR-ABL1-p190-3xHA-IRES-mCherry | This paper | N/A |
| MSCV-ABL1-3xHA-IRES-eGFP | This paper | N/A |
| MSCV-BCR-3xHA-IRES-tagBFP | This paper | N/A |
| MSCV-BCR-ABL1-p190 | A gift from Paul Ekert Lab | N/A |
| MSCV-SNX2-ABL1 | A gift from Paul Ekert Lab | N/A |
| MSCV-SFPQ-ABL1 | A gift from Paul Ekert Lab | N/A |
| **Cell line and bacteria strain** |  |  |
| HEK293 FT cell | American Type Culture Collection (ATCC) | CRL-3216 |
| Top10 chemically competent *E. coli* | Thermo Fisher | C404010 |
| Stbl3 chemically competent *E. coli* | Thermo Fisher | C737303 |
| **Tissue/bacteria culture reagents and materials** |  |  |
| Dulbecco's Modified Eagle's Medium (DMEM) Gibco | Thermo Fisher | 11965092 |
| GlutaMAX | Thermo Fisher | A1286001 |
| Opti-MEM I Reduced Serum Medium | Thermo Fisher | 11058021 |
| TrypLE Express Enzyme (1X) | Thermo Fisher | 12604013 |
| Heat-inactivated fetal Bovine Serum (FBS) | Thermo Fisher | 10100147 |
| LB broth | Thermo Fisher | 10855001 |
| Ampicillin | Sigma-Aldrich | A9393 |
| Penicillin/-Streptomycin | Thermo Fisher | 151401220 |
| Cell culture plasticwares | Greiner Bio-One, BD Biosciences, and Corning | NA |
| **Western blot antibodies** |  |  |
| $\beta$ actin monoclonal antibody (source: mouse application: 1:2000) | Sigma-Aldrich | A2228 |
| α-Tubulin Antibody (source: rabbit, application: 1:2000) | Sigma-Aldrich | SAB4500087 |
| Phospho-Stat5 (Tyr694) (D47E7) (source: rabbit, application 1:2000) | Cell signalling technology | 4322 |
| Stat5 (D2O6Y) (source: rabbit, application 1:2000) | Cell signalling technology | 94205 |
| P44/42 MAPK (Erk1/2) (source: rabbit, application 1:1000) | Cell signalling technology | 9102 |
| p44/42 MAPK (Erk1/2) P Thr202/Tyr204 (source: rabbit, application 1:1000) | Cell signalling technology | 9101 |
| (Horseradish peroxidase) HRP conjugated goat anti-mouse IgG secondary Antibody (application: 1:10,000) | Abcam | ab97023 |
| HRP conjugated goat anti-rabbit IgG secondary Antibody (application: 1:2000) | Abcam | ab205718 |
| IRDye® 680RD Goat anti-Rabbit IgG Secondary Antibody (0.1mg) (application: 1:10,000) | Li-cor | 92668071 |
| IRDye® 800CW Donkey anti-Mouse IgG Secondary Antibody (0.1mg) (application: 1:10,000) | Li-cor | 92632212 |
| **Cloning enzymes** |  |  |
| BbsI restriction enzyme | New England Biolabs | R3539 |
| NheI-HF restriction enzyme | New England Biolabs | R3131S |
| EcoRI restriction enzyme | Promega | R6011 |
| BamHI restriction enzyme | Promega | R6021 |
| T4 DNA ligase | Promega | M1801 |
| **DNA oligos for crRNAs cloning or gBLOCK for construct cloning** |  |  |
| crRNA sequences (supplementary Table 1) | Sigma or IDT | N/A |
| gBlock sequences (supplementary Table 2) | Sigma | N/A |
| Primer sequences (supplementary Table 3) | IDT | N/A |
| **Critical commercial assays or kits** |  |  |
| NucleoSpin Gel and PCR Clean-up Kit | Macherey-Nagel | 740609.50 |
| NucleoSpin Plasmid Mini kit for plasmid DNA | Macherey-Nagel | 740588.50 |
| **NucleoBond Xtra Maxi Plus** | Macherey-Nagel | 740416.50 |
| Lipofectamine 3000 transfection kit | Thermo Fisher | L3000015 |
| NucleoSpin RNA Plus | Macherey-Nagel | 740984.50 |
| Quick-RNA Miniprep Kit | Zymo Research | R1055 |
| high-capacity cDNA reverse transcription kit | Thermo Fisher | 4368814 |
| PowerUp™ SYBR™ Green Master Mix | Thermo Fisher | A25742 |
| Trans-Blot Turbo Midi 0.2 µm Nitrocellulose Transfer Packs | Bio-Rad | 1704159 |
| Phusion Site-Directed Mutagenesis Kit | Thermo Fisher | F541 |
| **Software and Algorithms** |  |  |
| Image J | National Institutes of Health | https://imagej.net |
| Prism software version 9 (GraphPad) | GraphPad Software Inc | https://www.graphpad.com/scientific-software/prism/ |
| Image Lab | Bio-Rad | https://www.bio-rad.com/en-au/product/image-lab-software?ID=KRE6P5E8Z |
| Image studio | Li-cor | https://www.licor.com/bio/image-studio-lite/ |
| RNAfold | Vienna RNAfold | http://rna.tbi.univie.ac.at/ |
| RNAplex | RNAplex | http://www.tbi.univie.ac.at/~htafer/ |
| IntaRNA | IntaRNA | http://rna.informatik.uni-freiburg.de/IntaRNA/Input.jsp;jsessionid=57BA904DBEE77BAA83BBAB1F20F55BAE |
| R package 'ggseqlogo' | CRAN-The Comprehensive R archive network | https://cran.r-project.org/web/packages/ggseqlogo/ |
| Matplotlib | Matplotlib | https://matplotlib.org/ |
| **Equipment** |  |  |
| EVOS M5000 FL Cell Imaging System | Thermo Fisher | AMF5000 |
| Trans-Blot Semi-Dry electrophoretic transfer cell | Bio-Rad | 1703940 |
| Trans-Blot Turbo Transfer System | Bio-Rad | 1704150 |
| Odyssey CLx Imager | Li-cor | 9140 |
| iBright CL1500 Imaging System | Thermo Fisher | A44114 |
| ChemiDoc Imaging System | Bio-Rad | 1708265 |
| StepOne Real-Time PCR system | Thermo Fisher | 4376357 |

## Notes

https://cas13b.github.io/

### Competing Interest Statement

The authors have declared no competing interest.

